# Structural mechanism of FusB-mediated rescue from fusidic acid inhibition of protein synthesis

**DOI:** 10.1101/2024.08.30.610483

**Authors:** Adrián González-López, Xueliang Ge, Daniel S.D. Larsson, Suparna Sanyal, Maria Selmer

**Affiliations:** Department of Cell and Molecular Biology, Uppsala University, BMC, P.O. Box 596, SE-75124 Uppsala, Sweden

**Keywords:** Fusidic acid, ribosome, EF-G, cryo-EM, elongation factor G, FusB

## Abstract

Antibiotic resistance protein FusB rescues protein synthesis from inhibition by fusidic acid (FA), which locks elongation factor G (EF-G) to the ribosome after GTP hydrolysis. Here, we present time-resolved single-particle cryo-EM structures explaining the mechanism of FusB-mediated rescue. FusB binds to the FA-trapped EF-G on the ribosome, causing large-scale conformational changes of EF-G that break ribosome interactions. This leads to dissociation of EF-G from the ribosome, followed by FA release. We also observe two independent binding sites of FusB on the classical-state ribosome, overlapping with the binding site of EF-G to each of the ribosomal subunits, yet not inhibiting tRNA delivery. Our results reveal an intricate resistance mechanism involving specific interactions of FusB with both EF-G and the ribosome, and a non-canonical release pathway of EF-G.

## INTRODUCTION

Antibiotic resistance is mediated by a range of different mechanisms, including drug efflux, mutations or modifications of the target, degradation or modification of the drug, and target protection^1^. Of these, target protection, involving a resistance protein that provides resistance through association with the antibiotic target, is the least characterized^2^, and the mechanism is unique for each target. One such example is FusB-type resistance to fusidic acid (FA)^3^.

FA is an antibiotic that primarily targets gram-positive bacteria, by inhibiting bacterial protein synthesis. It was introduced in the 1960s^4^ and is used topically against *Staphylococcus aureus* infections. FA binds to elongation factor G (EF-G), the GTPase translation factor involved in tRNA translocation^5^ and ribosome recycling^6^, preventing its release from the ribosome^7,8^. The binding site of FA is an interdomain pocket of EF-G lined by the sarcin-ricin loop (SRL) of 23S rRNA. This pocket is only formed in the ribosome-bound EF-G after GTP hydrolysis^9^, when switch I has transitioned to the GDP conformation. FA has been shown to stabilize switch II in its GTP conformation, inhibiting the conformational changes of EF-G required for release from the ribosome^9^. There are structures of FA-locked EF-G complexes in two ribosomal states: the chimeric state, where the head of the ribosomal small subunit (SSU) is swiveled and tRNA translocation is not completed^10^, and the post-translocational state^9^, where the ribosome and the tRNAs are in classical state.

The clinically most prevalent resistance mechanism to FA is FusB-type resistance, involving a resistance protein, FusB (or the homologs FusC, FusD or FusF), that provides low-level resistance to FA (minimum inhibitory concentration of 4-16 µg/ml, compared to 0.125 µg/ml for wild type)^11,12^ through binding to EF-G^13^. In addition, resistance can occur through mutations in EF-G^14^, through direct effects on FA binding, or through affecting EF-G stability or EF-G-ribosome interactions^15^. In laboratory experiments, mutations in uL6^16^, which interacts with EF-G^17^ can also provide resistance.

FusB is a two-domain protein, with an N-terminal alpha-helical bundle domain, and a C-terminal treble-clef zinc finger domain^18^. FusB binds with high affinity to *S. aureus* EF-G^19^, forming interactions with domains IV-V of EF-G^18,19^. Nuclear magnetic resonance (NMR) studies have shown that FusB binding causes increased dynamics of domain III of EF-G^20,21^. It has thus been proposed that FusB binding allosterically promotes disorder in domain III of EF-G, which causes release from the FA-inhibited ribosome^21^. FusB also increases the turnover of *S. aureus* EF-G on the ribosome in absence of FA^22^.

Here, we set out to determine the detailed structural mechanism of FusB-mediated rescue of FA inhibition. Using time-resolved cryo-EM, we solved multiple structures at different stages of rescue, including a pre-release complex which reveals how FusB releases EF-G from the ribosome and causes FA resistance. We additionally observe independent binding of FusB to the ribosomal A site, but our biochemical data shows no effect on aminoacyl-tRNA delivery. We describe an intricate mechanism of resistance that involves both FusB-mediated changes in the conformation of EF-G, and direct FusB-ribosome interactions.

## RESULTS

### Capturing FusB-mediated FA rescue

To capture FusB-mediated rescue using time-resolved cryo-EM, we prepared a FA-locked complex containing 70S ribosomes, a short synthetic mRNA, *E. coli* tRNA^fMet^, and *S. aureus* EF-G, in presence of FA and GTP. Initial tests with *S. aureus* 70S showed too low particle density for characterization of lowly populated states. Since FusB does not bind to *E. coli* EF-G^3^, we used the previously validated^18^ heterologous *E. coli* 70S-*S. aureus* EF-G complex for most of the work, but validated the highly populated states with *S. aureus* 70S. To allow the capture of short-lived rescue complexes on the cryo-EM grid with a standard plunging setup, the FA-locked ribosome complex was first incubated on the grid, followed by addition of FusB immediately before blotting and plunge-freezing. Initial tests indicated a diffusion-dependent gradient of ribosomal states, as two data sets collected on different parts of the same grid showed strikingly different populations of complexes (Extended data Figure 1). This method allowed grid vitrification 6 s after FusB addition, which is thus the upper limit for the reaction time of FA-inhibited complexes with FusB. An additional dataset was collected from an identical sample conventionally mixed in a tube 25 seconds before plunge freezing.

Through extensive focused 3D classification around the A-site region using a large number of classes, we identified six different complexes relevant to FA inhibition and rescue (Figure 1): FA-locked EF-G in chimeric state (CHI), FA-locked EF-G in post-translocational state (POST), FusB bound to EF-G in canonical POST state (FusB•EF-G•70S), FusB bound to domains IV-V of EF-G in a non-canonical ribosome binding site (FusB•EF-G•70S*), FusB bound to the decoding center of the SSU independently of EF-G (FusB•70S:SSU), and FusB bound to the large ribosomal subunit (LSU) and the P-site tRNA independently of EF-G (FusB•70S:LSU).

**Figure 1.**
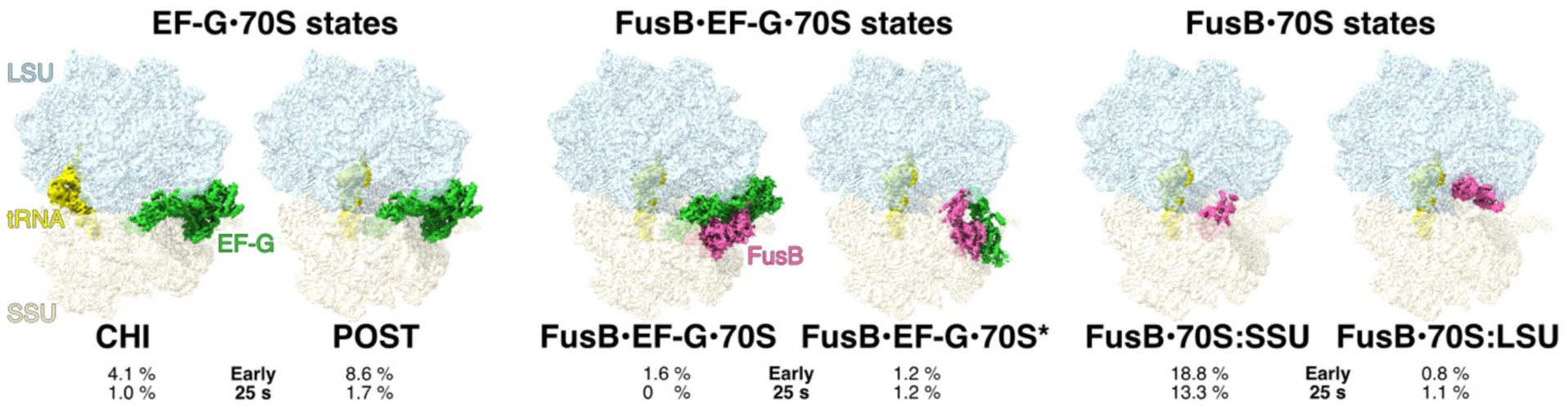
Time-resolved cryo-EM of FusB-mediated rescue of FA inhibition. Cryo-EM maps of six identified states are segmented 3 Å around the model and are showing the LSU (light blue), SSU (light yellow), tRNA (yellow), EF-G (green) and FusB (pink). The fraction of particles assigned to each state at each time point is indicated. Ribosome states with empty A-site, low-occupancy EF-G or A-site tRNA correspond to the remaining particle populations (64.9 % at the early time point and 81.7 % at the 25 s time point).

The global resolution of the maps ranges from 1.87 Å to 2.79 Å (Table 1), with local resolutions of EF-G and FusB ranging from 2.5 to 5 Å (Extended Data Figure 2).

**Table 1.** Cryo-EM refinement parameters and model validation.

|  | CHI<br>(EMDB-51350)<br>(PDB<br>9GHA) | POST<br>(EMDB-51351)<br>(PDB<br>9GHB) | FusB•<br>EF-G•70S<br>(EMDB-51352)<br>(PDB<br>9GHC) | FusB•<br>EF-G•70S*<br>(EMDB-51353)<br>(PDB<br>9GHD) | FusB•70S:<br>SSU<br>(EMDB-51354)<br>(PDB<br>9GHE) | FusB•70S:<br>LSU<br>(EMDB-51355)<br>(PDB<br>9GHF) | FusB•Sa70<br>S:SSU<br>(EMDB-51356)<br>(PDB<br>9GHG) | FusB•Sa70S:<br>LSU<br>(EMDB-51357)<br>(PDB<br>9GHH) |
| --- | --- | --- | --- | --- | --- | --- | --- | --- |
| **Data collection and processing** |  |  |  |  |  |  |  |  |
| Dataset | Early | Early | Early | 25 s | 25 s | 25 s | FusB•Sa70S | FusB•Sa70S |
| Magnification | 130 000 | 130 000 | 130 000 | 130 000 | 130 000 | 130 000 | 165 000 | 165 000 |
| Voltage (kV) | 300 | 300 | 300 | 300 | 300 | 300 | 300 | 300 |
| Electron exposure (e-/Å<sup>2</sup>) | 40 | 40 | 40 | 45.47 | 45.47 | 45.47 | 28.14 | 28.14 |
| Defocus range (µm) | -0.7 to -1.0 | -0.7 to -1.0 | -0.7 to -1.0 | -0.7 to -1.0 | -0.7 to -1.0 | -0.7 to -1.0 | -0.7 to -1.0 | -0.7 to -1.0 |
| Pixel size (Å) | 0.6480 | 0.6480 | 0.6480 | 0.6478 | 0.6478 | 0.6478 | 0.7280 | 0.7280 |
| Symmetry imposed | C1 | C1 | C1 | C1 | C1 | C1 | C1 | C1 |
| Initial particle images (no.) | 2 239 537 | 2 239 537 | 2 239 537 | 3 613 403 | 3 613 403 | 3 613 403 | 859 410 | 859 410 |
| Final particle images (no.) | 88 568 | 184 659 | 34 617 | 30 847 | 349 231 | 29 766 | 67 441 | 15 167 |
| Map resolution (Å) | 2.24 | 2.21 | 2.79 | 2.41 | 1.87 | 2.40 | 2.22 | 2.7 |
| FSC threshold | 0.143 | 0.143 | 0.143 | 0.143 | 0.143 | 0.143 | 0.143 | 0.143 |
| **Refinement** |  |  |  |  |  |  |  |  |
| Initial model used (PDB code) | 9GHE | 9GHE | 9GHE | 9GHE | 8CGK,<br>8CGJ,<br>8CF1,<br>7K00,<br>7N2C,<br>4ADN | 9GHE | 8P2F,<br>4ADN | 9GHG |
| Model resolution (Å) | 2.2 | 2.2 | 2.8 | 2.4 | 1.9 | 2.4 | 2.2 | 2.7 |
| FSC threshold | 0.5 | 0.5 | 0.5 | 0.5 | 0.5 | 0.5 | 0.5 | 0.5 |
| Post-processing | Local<br>filtering | Local<br>filtering | Local<br>filtering | Local<br>filtering | Local<br>filtering | Local<br>filtering | Local<br>filtering | Local<br>filtering |
| Map sharpening B factor. (Å<sup>2</sup>) | -35 | -36 | -25 | -33 | -36 | -33 | -32 | -23 |
| **Model composition** |  |  |  |  |  |  |  |  |
| Non-hydrogen atoms | 145 564 | 145 490 | 147 172 | 143 666 | 146 257 | 141 978 | 150 404 | 144 758 |
| Protein residues | 6 176 | 6 188 | 6 380 | 5 946 | 5 718 | 5 718 | 5 685 | 5 680 |
| RNA residues | 4 510 | 4 499 | 4 502 | 4 498 | 4 501 | 4 501 | 4 654 | 4 650 |
| Waters | 0 | 0 | 0 | 0 | 4 235 | 0 | 5 514 | 0 |
| Magnesium atoms | 272 | 303 | 301 | 303 | 303 | 303 | 145 | 145 |
| **B factors (Å<sup>2</sup>)** |  |  |  |  |  |  |  |  |
| Protein | 72 | 71 | 81 | 69 | 57 | 62 | 72 | 91 |
| RNA | 57 | 54 | 62 | 51 | 45 | 50 | 65 | 79 |
| Ligand | 41 | 44 | 49 | 39 | 30 | 38 | 30 | 44 |
| Water | - | - | - | - | 24 | - | 38 | - |
| **R.m.s. deviations** |  |  |  |  |  |  |  |  |
| Bond lengths (Å) | 0.008 | 0.008 | 0.008 | 0.008 | 0.008 | 0.008 | 0.008 | 0.008 |
| Bond angles (°) | 1.187 | 1.183 | 1.175 | 1.191 | 1.294 | 1.195 | 1.193 | 1.198 |
| **Validation** |  |  |  |  |  |  |  |  |
| MolProbity score | 1.30 | 1.12 | 1.22 | 1.06 | 1.04 | 1.12 | 1.38 | 1.38 |
| Clashscore | 3.47 | 2.87 | 3.60 | 2.5 | 2.54 | 3.02 | 4.17 | 3.49 |
| Poor rotamers (%) | 1.24 | 0.83 | 1.23 | 1.08 | 0.95 | 1.10 | 1.73 | 1.69 |
| **Ramachandran plot** |  |  |  |  |  |  |  |  |
| Favored (%) | 97.57 | 97.79 | 98.50 | 98.66 | 98.52 | 98.68 | 98.67 | 98.78 |
| Allowed (%) | 2.31 | 2.08 | 1.44 | 1.29 | 1.45 | 1.29 | 1.31 | 1.20 |
| Disallowed (%) | 0.12 | 0.13 | 0.06 | 0.05 | 0.04 | 0.04 | 0.02 | 0.02 |
| Rama-Z score | -0.54 | -0.54 | 1.19 | 0.88 | 1.13 | 1.23 | 1.07 | 0.99 |

The complexes show different abundance at the two time points, demonstrating FusB-mediated rescue of FA inhibition (Figure 1). The pre-rescue FA-locked EF-G complexes (CHI and POST) dramatically decrease in population between the early and late time points from 12.7% to 2.7% of the total ribosome population, respectively. The canonical rescue complex FusB•EF-G•70S is only present in the early dataset and at a relatively low proportion (1.6% of the total ribosomes), indicating that it is an early and transient complex and that most ribosomes have already been rescued at this time point. The other FusB·EF-G complex (FusB•EF-G•70S*) is present in both datasets at 1.2%. Strikingly, the FusB•70S:SSU complex constitutes 19% and 13% of the ribosomes at the two time points, showing that FusB has a strong propensity for ribosome binding also independently of EF-G.

### Locking of *S. aureus* EF-G to *E. coli* 70S by FA

The CHI and POST structures show that *S. aureus* EF-G is locked in a close to identical structure on the *E. coli* ribosome as on the *S. aureus* ribosome^17^ (Supplementary Figure 1a-b). As expected, EF-G binds in an extended conformation to the ribosome, spanning from the SSU A-site to the SRL of the LSU. FA is bound in its pocket next to GDP and a magnesium ion (Supplementary Figure 1c).

### FusB•EF-G on the ribosome

In the early rescue complex, FusB•EF-G•70S, FusB is bound to the POST-state EF-G (Figure 2a), and there is clear density showing that FA and GDP remain bound to EF-G (Supplementary Figure 2d-g). In this state, FusB makes extensive contact with EF-G through both of its domains (Figure 2b). The N-terminal domain of FusB is wedged between domains I and V of EF-G and also interacts with domains II-III of EF-G. The C-terminal domain of FusB forms an extended β-sheet with domain IV of EF-G and also interacts with domains III and V.

**Figure 2.**
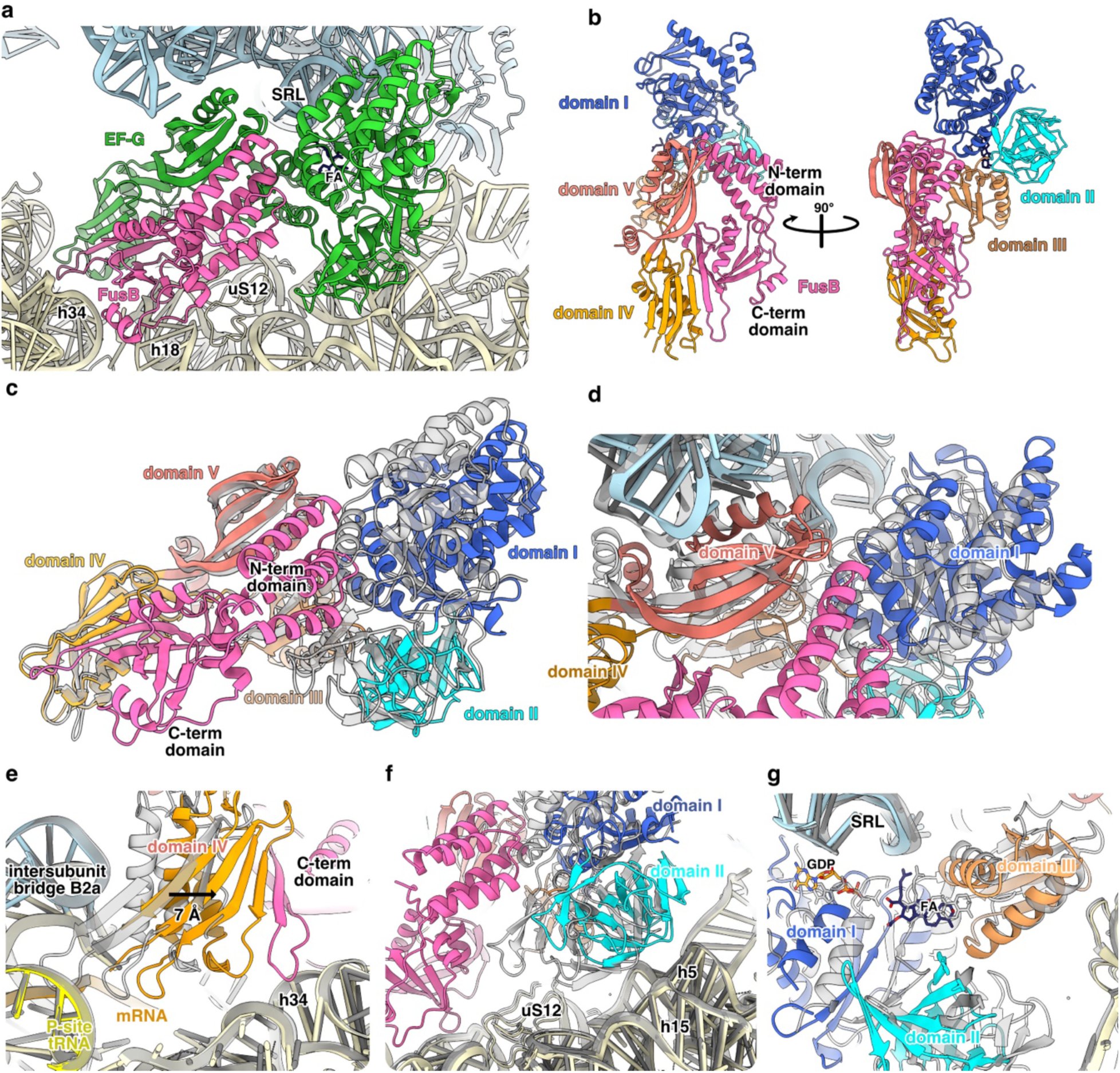
Structure of the FusB•EF-G•70S FA rescue complex. **(a)** Overall structure of the FusB•EF-G•70S complex. **(b)** Overview of the FusB•EF-G complex in FusB•EF-G•70S structure. EF-G is colored by domain. **(c)** Comparison of EF-G in FusB•EF-G•70S (colors as in b) with EF-G in the POST conformation (gray) when domains IV-V of EF-G are aligned. **(d-g)** Movement of EF-G domains from the POST state (gray) to the FusB•EF-G•70S complex with structures aligned by 23S rRNA (colors as in b, tRNA in yellow and mRNA in orange).

Alignment of the FusB•EF-G•70S and POST complex structures shows that FusB causes a major conformational change of EF-G (Figure 2c-g, Supplementary Figure 2a-d, Supplementary Video 1). The N-terminal domain of FusB would otherwise clash with helix 3 of domain I of EF-G (Figure 2c). Instead, it pushes domains I-III away from domains IV-V (Figure 2d). Meanwhile, the C-terminal domain of FusB is anchored to the SSU and pulls domain IV away from the tRNA and mRNA (Figure 2e), with a 7 Å shift of the beta-sheet of domain IV. The interactions between FusB and the ribosome mainly occur between two loops (K99-K105, K174-S177) and helices h18 and h34. The interactions with h34 seem to provide specificity for the POST-state ribosome, since this helix is part of the SSU head that moves between the CHI and POST states.

Alignment of the FusB•EF-G•70S and POST complex structures based on 23S rRNA shows that all domains of EF-G are subject to conformational change (Figure 2d-g, Supplementary Figure 2a-d), while the ribosome is unaffected. This results in an overall loss of interactions between EF-G and the ribosome (the number of EF-G residues within 4 Å of the ribosome decreases from 41 to 23). The structural integrity of the FA-binding pocket and density for FA (Supplementary Figure 2e-g) shows that FusB primarily promotes EF-G dissociation from the ribosome, which, in turn, leads to FA dissociation since FA has very low affinity to free EF-G^23^.

### FusB binds 70S ribosomes independently of EF-G

Surprisingly, FusB also binds to two different sites of the ribosome without EF-G (Figure 3, Supplementary Figure 3). In the most populated complex (FusB•70S:SSU), FusB is bound at the decoding center of the SSU of the classical-state ribosome. The C-terminal domain of FusB makes direct contact with the monitoring bases A1492 and A1493 in h44 (Figure 3b). The side chains of the lysine-rich loop of FusB interact with h31, h32 and h34 (Figure 3c). FusB also interacts with h18, the mRNA, uS12, and the intersubunit bridge B2a.

**Figure 3.**
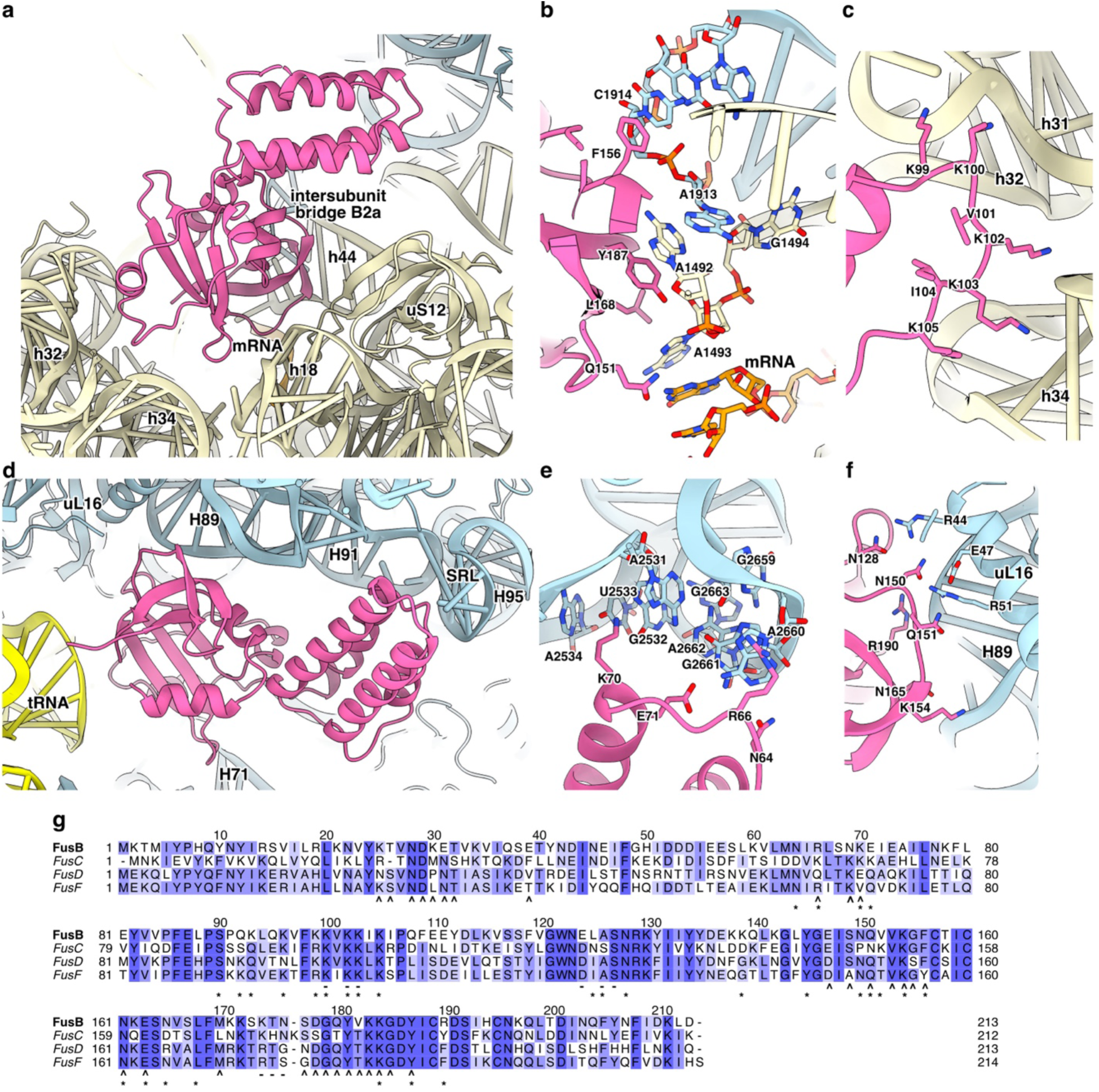
Independent binding of FusB to the ribosome. (a) 30S binding site of FusB (FusB•70S:SSU). (b) FusB interaction with intersubunit bridge B2a. (c) The lysine-rich loop of FusB contacts h31, h32 and h34 of 16S rRNA. (d) 50S binding site of FusB (FusB•70S:LSU). (e) Interaction of FusB with H91 and the SRL of 23S rRNA. (f) FusB interaction with H89 and uL16. (g) ClustalO multiple sequence alignment of FusB (Uniprot Q8GNY5), FusC (WP_001033157), FusD (WP_011303797) and FusF (BAQ33930). Residues that interact with the ribosome in FusB•70S complexes (*), and with EF-G (^) or the ribosome (-) in FusB•EF-G•70S are marked below the alignment.

In the FusB•70S:LSU complex, FusB contacts the LSU at H71, H89, H91, the SRL at H95, uL16, and the P-site tRNA of the classical-state ribosome (Figure 3d-f). The region of the C-terminal domain of FusB close to the tRNA, including the lysine-rich loop, is more disordered than the rest of FusB.

FusB would in both of its independent binding sites sterically clash with EF-G in the FA-locked state, showing that FusB directly competes with EF-G to interact with the ribosome. Multiple sequence alignment (Figure 3g) shows that most of the ribosome-interacting residues are conserved also in FusC, FusD, and FusF, suggesting that the other variants can bind to the ribosome in the same way. The residues involved in interactions with EF-G are similarly to a large extent conserved. Remarkably, there is a large overlap between residues in FusB that interact with the ribosome (in the FusB•70S complexes), and with EF-G (in FusB•EF-G•70S and FusB•EF-G•70S*). For example, F156 and Y187, which, if mutated to alanine, make FusB unable to bind to EF-G^19^, interact directly with C1914 from 23S rRNA (Figure 3b) and A1492 from 16S rRNA, respectively. Hence, binding of FusB to EF-G and to its independent sites on the ribosome are mutually exclusive.

To determine if the observed binding of FusB to the ribosome would impair tRNA delivery, dipeptide and tripeptide formation assays were performed using a single-turnover, fast-kinetics approach. Here, 70S initiation complex (IC) was pre-incubated with an excess of FusB and subsequently combined with elongation mixture (EM) in a quench-flow instrument. The presence of FusB in the IC had no effect on dipeptide formation, the rate remaining nearly identical, at 36.6 ± 1.2 s⁻¹ without FusB and 35.6 ± 1.3 s⁻¹ with FusB (Figure 4a). This indicates that FusB binding to the ribosome does not inhibit EF-Tu-mediated tRNA delivery. FusB, however, did decrease the tripeptide formation rate by approximately 2.5-fold when *S. aureus* EF-G was used, from 0.54 ± 0.01 s⁻¹ in the absence of FusB to 0.22 ± 0.09 s⁻¹ when FusB was present (Figure 4b). After dipeptide formation, both the independent binding sites of FusB would be sterically blocked by the A-site tRNA. Thus, inhibition of tripeptide formation is likely due to sequestering of EF-G, since FusB has high affinity to *S. aureus* EF-G in solution^22^. This explanation is further supported by a control tripeptide formation experiment with *E. coli* EF-G, to which FusB does not bind^3^, showing no inhibitory effect (Figure 4c).

**Figure 4.**
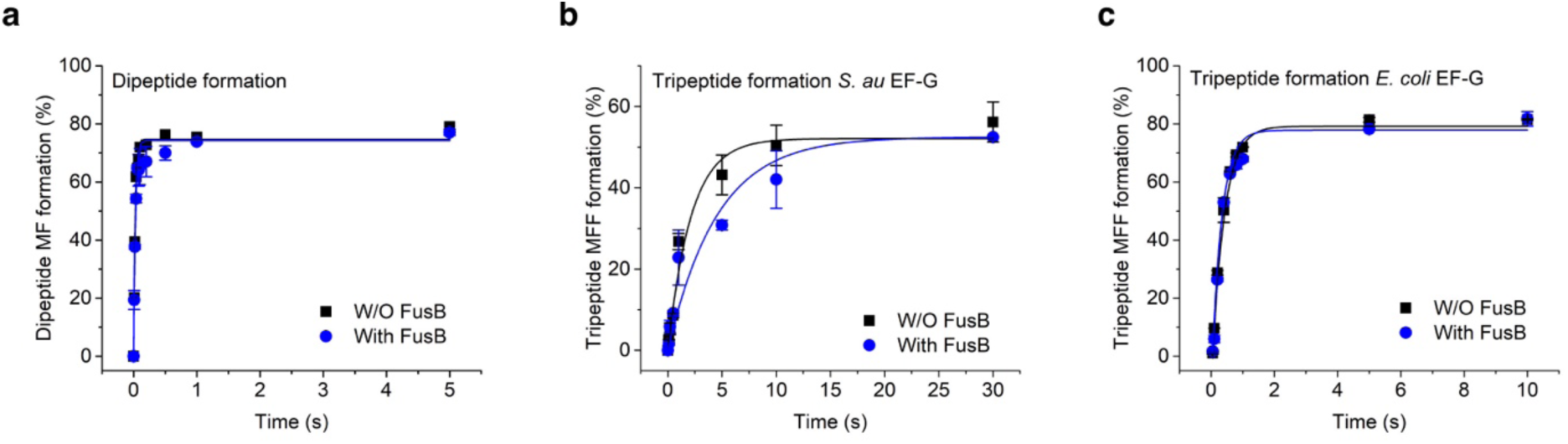
Di- and tripeptide formation experiments in absence (black) and presence (blue) of FusB in two experimental replicates showing the mean and standard deviation. **(a)** Comparison of single-turnover dipeptide formation with *S. aureus* EF-G. **(b)** Comparison of single-turnover tripeptide formation with *S. aureus* EF-G. **(c)** Comparison of single-turnover tripeptide formation with *E. coli* EF-G.

### FusB binds identically to both *E. coli* and *S. aureus* ribosomes

To further validate the observed binding of FusB to the ribosome, a control sample was prepared with *S. aureus* 70S ribosomes, *E. coli* tRNA^fMet^, a short synthetic mRNA, and FusB. As expected, FusB formed the same two complexes with *S. aureus* 70S: FusB•Sa70S:SSU and FusB•Sa70S:LSU. In both cases, FusB and its interactions with the ribosome are close to identical to the heterologous complexes (Supplementary Figure 4). This is not surprising, as all the interacting nucleotides and amino acid residues are conserved between *E. coli* and *S. aureus*, except for R51 (I52 in *S. aureus*) in uL16. In this dataset, FusB was found in 12.3 % of the total ribosomes (10 % FusB•Sa70S:SSU and 2.3 % FusB•Sa70S:LSU).

### FusB•EF-G binds to the ribosome outside of the A-site

One of the identified complexes (FusB•EF-G•70S*) shows FusB bound to domains IV-V of EF-G, with no visible density for domains I-III of EF-G (Figure 1). In this state, domain IV of EF-G does not contact the ribosome (Figure 5a). EF-G interacts with the SRL, the stem-loops of H43 and H44, and uL6. FusB contacts h18 (through its lysine-rich loop), h34, the stem-loops of H43 and H89, and uS12. The FusB-EF-G interface is the same way as in the early complex (Figure 5b), apart from interactions with the disordered domains I-III of EF-G. If these would retain the same conformation as in FusB•EF-G•70S, they would clash with the LSU. Furthermore, the conformations of these domains in crystal structures of EF-G would also clash with FusB (Figure S5).

**Figure 5.**
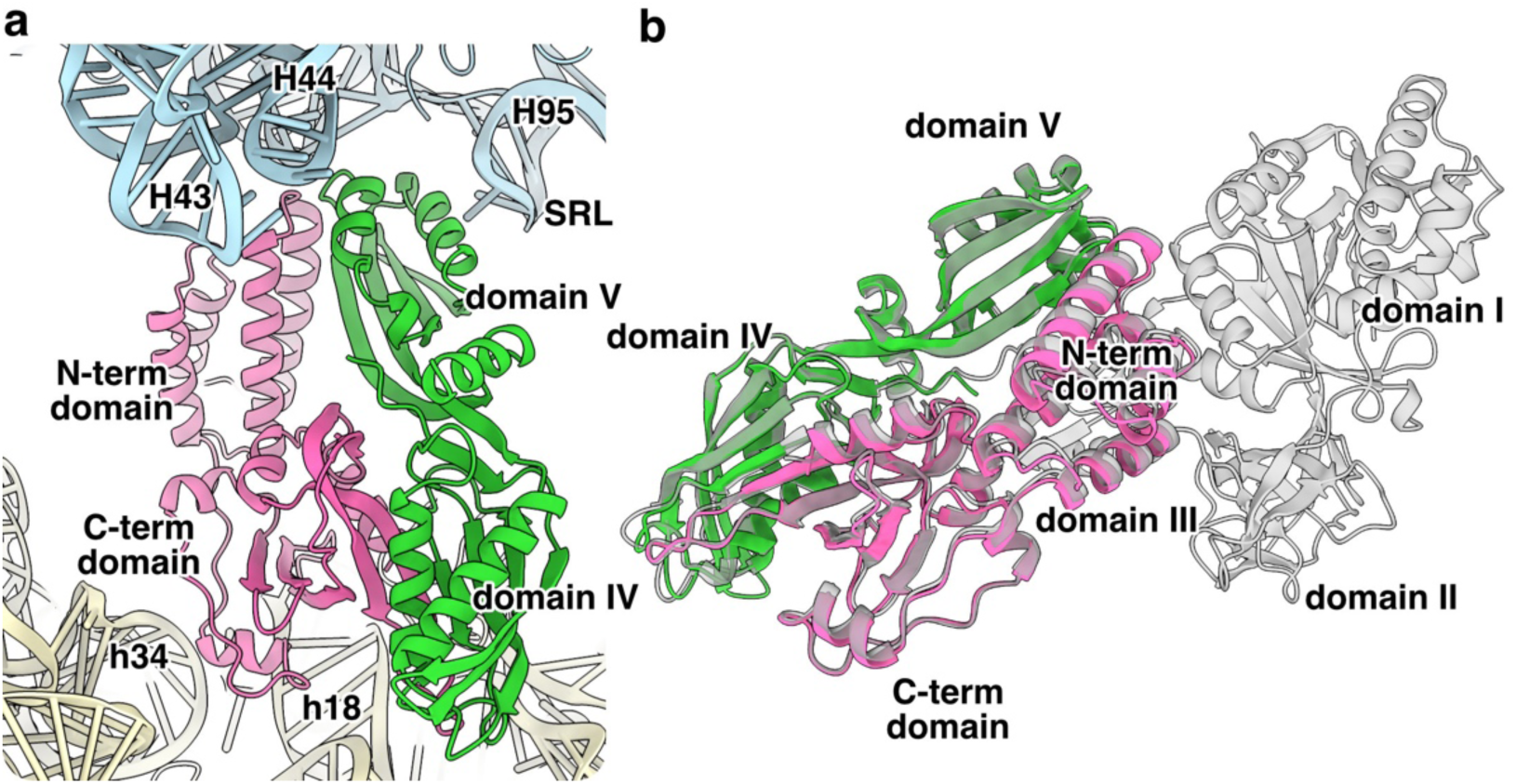
(a) Non-canonical ribosome binding of FusB•EF-G in FusB•EF-G•70S*. (b) Comparison of the FusB-EF-G complex in FusB•EF-G•70S* (magenta and green) and FusB•EF-G•70S (gray). The structures are aligned to EF-G domains IV-V. The RMSD is 0.589 Å over 204 Cα atoms for EF-G, and 0.582 Å over 212 Cα atoms for FusB.

The orientation of this complex differs by almost 180° compared to the early complex. Therefore, it is unlikely to represent a late rescue intermediate as it would require a large rotation of the early complex, while keeping domain V of EF-G between H44 and H95. Since the proportion of FusB•EF-G•70S* is the same at both time points, this more likely represents a rebinding event of FusB•EF-G.

## DISCUSSION

In this work, we have solved the structure of a short-lived antibiotic resistance rescue complex of FusB bound to the FA-inhibited ribosome. This state could not be observed at 25 s after reaction start, which is the shortest time point we could achieve by conventional plunge-freezing. Instead, we took advantage of the slow on-grid diffusion of FusB, which effectively created a concentration distribution, and thus a reaction time distribution of ribosomal complexes on the grid. The majority of FA-inhibited ribosomes had already been rescued, but this simple methodology in combination with advanced particle classification gave access to high-resolution information of an otherwise non-observable state in this process. We envision that this procedure will also be applicable and provide insights into other biological processes, since it offers a readily accessible approach in absence of advanced instrumentation for time-resolved grid preparation.

Using this methodology, we obtained structures of six different complexes related to FusB-mediated rescue of FA inhibition. Based on these, we propose a detailed mechanism of FusB-type resistance (Figure 6). FA locks EF-G to the ribosome in two possibly interchanging states (CHI and POST structures). FusB binds to EF-G in the POST state (FusB•EF-G•70S). Here, FusB induces a major conformational rearrangement of EF-G by wedging its N-terminal domain between domains I and V and by interacting with both the SSU and domain IV of EF-G. This leads to an overall loss of contacts between EF-G and the ribosome, causing release of the FusB•EF-G complex from the ribosome, and subsequent dissociation of FA from EF-G. Other FusB molecules can bind to either subunit of the classical-state 70S ribosome, and both would sterically clash with EF-G in the FA-locked conformation. This suggests that FusB may prevent the previously observed re-binding of EF-G•FA^8^, while maintaining the ribosome in a state that is capable of continuing translation by binding of a new aminoacyl tRNA-EF-Tu ternary complex.

**Figure 6.**
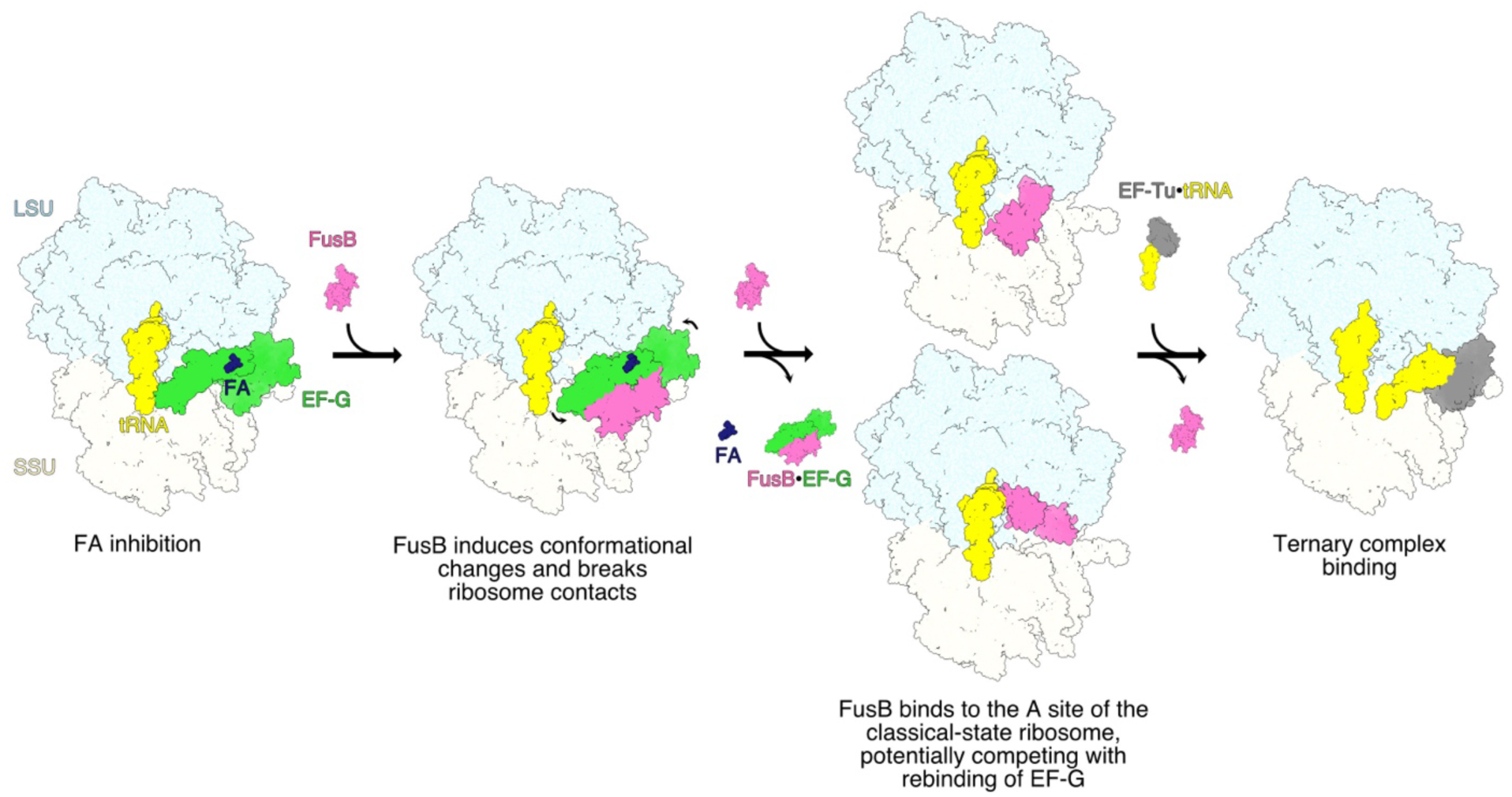
Proposed mechanism of FusB-mediated resistance.

During the canonical translocation cycle of EF-G, GTP hydrolysis is followed by conformational changes of the switch regions of domain I, inducing inter-domain conformational changes required for dissociation from the ribosome^24^. Time-resolved cryo-EM studies of translocation indicate that domains I-III of EF-G dissociate first from the ribosome, while domain IV retains its normal position until release^25^. Binding of FusB was previously proposed to cause release of the FA-inhibited state “by allosteric effects on the dynamics of the drug target” that would allow transmission of conformational change from the GTPase domain to domain IV-V of EF-G. Here, we have instead shown that EF-G is released in presence of FA through a new conformational pathway. Our structures suggest that the increased dynamics of domain III observed in the free FusB-EF-G complex^21^ are a consequence of lost contacts with the ribosome in combination with steric clashes of FusB with interdomain contacts observed in the crystal structures of EF-G (Supplementary Figure 5). Domain III appears to be the most dynamic of the domains in EF-G, since it displays disorder in many structures of free EF-G^15,26^. The observed steric block likely destabilizes this domain further, uncoupling domains I-II from domains IV-V of EF-G, explaining why only domains IV and V of EF-G are ordered relative to the ribosome in the non-canonical FusB•EF-G•70S* complex.

FA resistance mutations in EF-G (FusA) have been described to interfere with direct FA interactions, EF-G-ribosome interactions, the structural stability of EF-G and interdomain contacts in EF-G^15^. Our results show that FusB-mediated resistance similarly involves alteration of all these properties apart from the direct FA-EF-G interactions.

Comparison of our FusB•EF-G structures on the ribosome (FusB•EF-G•70S and FusB•EF-G•70S*) with a published NMR-based model of a truncated version of the free complex lacking domains I-II of EF-G, shows large discrepancies (Supplementary Figure 6), for example in the interaction between the C-terminal domain of FusB and domain IV of EF-G. We find it likely that the FusB-EF-G interface observed in our structures will be retained in solution.

Remarkably, the FusB•70S complexes are at both time points more abundant than EF-G•70S complexes in our sample conditions. Even though FusB would directly clash with an A-site tRNA, it does not inhibit dipeptide formation (Figure 3h), even at high concentration. However, in these conditions, FusB inhibits EF-G function, as tripeptide formation kinetics show a two-fold rate reduction. This is likely due to the formation of FusB•EF-G complex before translocation due to their high affinity^19^. Thus, the slower translocation is either due to a reduced concentration of free EF-G or indicative that the FusB-EF-G complex can translocate at a slower rate. The direct binding of FusB to the ribosome may also have some other function in translation, unrelated to FA resistance, since homologs of FusB are found in the chromosome of gram-positive bacteria that are unlikely to have encountered the antibiotic^3^.

Production of FusB on the clinically isolated pUB101 plasmid is regulated by translational attenuation^3^, in line with a requirement for carefully controlled concentration of FusB to provide rescue and not inhibition at the physiological concentrations of the components of the translation machinery. We expect that FusB *in vivo* is sub-stoichiometric to EF-G to avoid inhibition of translocation, whereas our biochemical experiments have FusB in excess to EF-G.

In contrast to other target protection mechanisms of resistance, such as tetracycline ribosomal protection proteins and ABC-F-mediated resistance (reviewed in^2^), FusB does not interact directly with the antibiotic, and its action does not induce the direct release of the antibiotic. Instead, the target protein (EF-G) is released from the ribosome, which in turn leads to the release of the antibiotic, as FA does not bind to EF-G in solution^23^. Unfortunately, this makes development of FA analogs or derivatives that overcome FusB-type resistance challenging, and other strategies may be needed.

In this study, we have elucidated the structural mechanism of FusB-mediated rescue of FA inhibition, but some details still remain unclear. Future studies will clarify the interplay of FusB and EF-G *in vivo*, and whether the ribosome binding of FusB or its homologs has additional functions in bacterial protein synthesis.

## MATERIALS AND METHODS

### Cloning, overexpression and purification of *S. aureus* FusB and EF-G

*S. aureus* EF-G was purified as published^17^ by immobilized metal chromatography and size exclusion chromatography. The final protein sample was stored in 50 mM Tris-HCl pH 7.5, 300 mM NaCl, 5 mM β-mercaptoethanol at -70 °C. This construct encodes a 6xHis tag with a TEV protease cleavage site and a linker to the EF-G sequence. After TEV cleavage, only Ser-Leu remains at the N-terminus.

The *S. aureus fusB* gene was cloned into pEXP5-NT-TOPO (Invitrogen) following the manufacturer’s instructions (pEXP5-NT_FusB). The sequence was confirmed by Sanger sequencing. This construct encodes a 6xHis tag with a TEV protease cleavage site and a linker to the FusB sequence. After TEV cleavage, only Ser-Leu remains at the N-terminus.

pEXP5-NT_FusB was transformed into *Escherichia coli* BL21 (DE3). An overnight culture was inoculated 1:100 into a 2.8 l baffled flask with 800 ml of LB with 100 µg/ml of ampicillin.

Protein expression was induced at an OD600 nm of 0.5-0.6 with 1 mM isopropyl-β-thiogalactopyranoside and the culture was incubated for 16-18 h at 30 °C. The cells were harvested at 7 400 x g for 30 min in a JLA 9.1000 rotor (Beckman Coulter, Brea, California, USA), washed with 150 mM NaCl, and stored at -20 °C.

The cells were resuspended in lysis buffer (50 mM Tris-HCl pH 7.5, 300 mM NaCl, 20 mM imidazole) with 0.1 % (v/v) Triton X-100, one EDTA-free mini-Complete protease inhibitor tablet (Roche, Basel, Switzerland) and DNase I. Then, they were lysed in a flow cell disruptor (Constant Systems Ltd., Daventry, United Kingdom). The lysate was centrifuged at 27 000 x g for 45 min in an SS-34 rotor (Thermo Fisher Scientific, Waltham, MA, USA).

The supernatant was filtered by 0.45 µm with a polyethersulfone syringe filter (Sarstedt AG & Co, Nümbrecht, Germany) and incubated with 1 ml of Ni Sepharose Fast Flow (Cytiva, Uppsala, Sweden) equilibrated with lysis buffer. The column was washed with 25 column volumes of 600 mM NaCl in lysis buffer at pH 7.8 and the protein was eluted with 400 mM imidazole in lysis buffer at pH 7.8. The eluate was exchanged into GF buffer (20 mM Tris-HCl pH 7.8, 300 mM NaCl, 5 mM β-mercaptoethanol) using a PD-10 column (Cytiva, Uppsala, Sweden). The 6x-histidine-tag was removed by cleavage with 1:25 molar ratio of TEV protease at 8 °C for 16-18 h followed by reverse immobilized affinity chromatography on 1 ml of Ni Sepharose Fast Flow.

FusB was purified further by gel filtration on a Hiload 16/60 Superdex-75 column (GE Healthcare, Uppsala, Sweden) equilibrated with GF buffer. The peak fractions were concentrated to 5-10 mg/ml with a 10-kDa cutoff Vivaspin Turbo 15 (Sartorius AG, Göttingen, Germany), frozen in liquid nitrogen, and stored at -70 °C.

### Preparation of components for cryo-EM

*E. coli* 70S ribosomes were purified from *E. coli* MRE600 cells according to^27^. *S. aureus* NCTC 8325-4 ribosomes were purified as published^17^. The final 70S sample was stored in HEPES polymix buffer (5 mM HEPES pH 7.5, 5 mM NH4Cl, 5 mM Mg(OAc)2, 100 mM KCl, 0.5 mM CaCl2, 8 mM putrescine, 1 mM spermidine, and 1 mM dithioerythritol) at −70 °C. mRNA Z4AUGGCA (5’-GGCAAGGAGGUAAAAAUGGCAAAA-3’) was produced by chemical synthesis (Dharmacon) and *E. coli* tRNA^fMet^ was overexpressed and purified as published^28^.

### Di- and tri-peptide formation assay

The translation components used in the biochemical assays are from *E. coli* unless mentioned otherwise. His-tagged initiation factors (IF1, IF2 and IF3), elongation factors (EF-Tu, EF-Ts and EF-G), 70S ribosome, XR7 mRNA with the coding sequence Met-Phe-Phe-Stop (AUG-UUC-UUU-UAA) and f[^3^H]Met-tRNA^fMet^ were prepared according to^29,30^. The 70S initiation complex (IC) and elongation mixture (EM) were prepared separately by incubating the respective components in HEPES polymix (pH 7.5) buffer^29^ at 37 °C for 15 minutes. The IC was assembled by combining 1 μM 70S ribosomes, 10 μM XR7 MFF mRNA, 1 μM f[³H]Met-tRNAᶠᴹᵉᵗ, and 2 μM each of the three initiation factors (IF1, IF2, and IF3). 10 μM *S. aureus* FusB was added to IM to test its effect on di and tripeptide formation. The EM was prepared by mixing 10 μM EF-Tu, 10 μM EF-Ts, 10 μM tRNA^P^ʰᵉ, 0.2 mM Phe amino acid, 1 unit of Phe tRNA synthetase, and 2.5 μM *S. aureus* or *E. coli* EF-G (as indicated). Both mixtures were supplemented with an energy regeneration system comprising 1 mM GTP, 1 mM ATP, 10 mM phosphoenolpyruvate, 0.05 mg/mL pyruvate kinase, and 0.002 mg/mL myokinase. To capture the kinetics of peptide formation, equal volumes of the IM and EM were rapidly mixed in a Quench-Flow instrument (RQF-3, KinTek Corp.). The reactions were quenched by adding 17% formic acid at specific time points. Following quenching, peptides were released by KOH treatment. Mono-, di- and tri-peptides were separated using a C18 reverse-phase column connected to a high-performance liquid chromatography system and detected with a β-RAM radioactive detector^29^. The di and tripeptide formed was plotted against time and the rate of di / tripeptide formation was determined by fitting the data to a single exponential curve using Origin Pro 2016 software. This was done with two distinct experimental replicates (Supplementary Figure 7) from which the means and standard deviations were calculated.

### EF-G•Ec70S complex preparation for grid vitrification

The sample was prepared by mixing 0.75 µM (final concentrations) 70S *E. coli* ribosomes in HEPES polymix buffer (with 20 mM HEPES pH 7.5 and 5 mM BME as reducing agent) with 3 µM mRNA Z4AUGGCA and incubated for 10 min at 37 °C. Then, 3 µM *E. coli* tRNA^fMet^ was added and the mix was incubated for 10 min at 37°C. Next, 7.5 µM *S. aureus* EF-G, 400 µM FA (Sigma-Aldrich, Merck, Darmstadt, Germany), and 1.5 mM GTP were added, followed by incubation for 10 min at 37 °C, then kept on ice until plunge-freezing.

### FusB•*Sa*70S complex preparation for grid vitrification

The sample was prepared by mixing 0.5 µM (final concentrations) 70S *S. aureus* ribosomes in HEPES polymix buffer with 5 µM mRNA Z4AUGGCA and incubated for 10 min at 37 °C. Then, 5 µM *E. coli* tRNA^fMet^ was added and the mix was incubated for 10 min at 37°C. Less than 30 s before plunge-freezing, 15 µM FusB was added.

### Cryo-EM grid preparation

All grids were prepared in a similar way with minor differences.

#### Early sample

A QuantiFoil 200-mesh R 2/1 grid with 2 nm continuous carbon (QuantiFoil Micro Tools GmbH, Großlöbichau, Germany) was glow-discharged 15 s at 20 mA and 0.39 mBar using EasiGlow (Ted Pella, Inc., Redding, CA, USA). 3 µl of EF-G•*Ec*70S was incubated on the grid for 30 s at 4 °C and 95 % humidity. Then, 1.2 µl of FusB 18.5 µM in HEPES polymix buffer with 400 µM FA was added into the drop, blotted for 3 s, and plunge-frozen into liquid ethane in a Vitrobot Mark IV (Thermo Fisher Scientific, Waltham, MA, USA) at 4 °C and 95 % humidity. The total time from adding FusB and the grid touching the ethane was 6 s.

#### 25 s sample

2 µl of 18.5 µM FusB was mixed with 5 µl of EF-G•*Ec*70S on ice and directly added to a glow-discharged (same as the early sample) QuantiFoil 200-mesh R 2/1 grid with 2 nm continuous carbon, blotted for 4 s and plunge-frozen in a Vitrobot Mark IV. The total time between adding FusB and the grid touching the ethane was 25 s.

#### FusB•Sa70S sample

A QuantiFoil 300-mesh R 2/2 grid with 2 nm continuous carbon was glow-discharged for 30 s. Then, 3 µl of FusB•*Sa*70S sample was added to the grid, incubated for 10 s, blotted for 3 s and plunge-frozen in a Vitrobot Mark IV.

#### Preliminary dataset

A QuantiFoil 300-mesh R 2/2 grid with 2 nm continuous carbon was glow-discharged for 30 s. 2.5 µl of EF-G•*Ec*70S (0.25 µM 70S ribosomes, 1 µM mRNA, 1 µM tRNA^fMet^, 2.5 µM EF-G, 400 µM FA, and 0.5 mM GTP) was incubated on the grid for 30 s.

Then, 1 µl of FusB 12.5 µM in HEPES polymix buffer with 400 µM FA was mixed into the drop by pipetting, blotted for 4 s, and plunge-frozen in a Vitrobot Mark IV at 4 °C and 95 % humidity. The total time from adding FusB and the grid touching the ethane was 10 s.

### Cryo-EM data collection

All the grids were screened on a Glacios TEM operated at 200 kV equipped with a Falcon-III direct electron detector (Thermo Fisher Scientific, Waltham, MA, USA).

#### Early dataset

Collected on a Titan Krios G3i (Thermo Fisher Scientific, Waltham, MA, USA) operated at 300 kV and equipped with a K3 BioQuantum direct electron detector (Gatan, Inc, AMETEK, Berwyn, PA, USA) and energy filter using 20 eV slit. The data were acquired at 130 000 x nominal magnification with a calibrated pixel size of 0.648 Å. A total of 24 479 movies were collected in 40 frames with a total dose of 40 e^-^/Å2 (16.8 e^-^/pixel/s) over 1 s with a set defocus between -0.7 to -1.0 µm.

#### 25 s dataset

Collected on a Titan Krios operated at 300 kV and equipped with a K3 BioQuantum direct electron detector and energy filter using 20 eV slit. The data were acquired at 130 000 x nominal magnification with a calibrated pixel size of 0.6478 Å. A total of 32 937 were collected in 45 frames with a total dose of 45.47 e^-^/Å2 (15.9 e^-^/pixel/s) over 1.2 s with a set defocus between -0.7 to -1.0 µm.

#### FusB•Sa70S dataset

Collected on a Titan Krios G2 operated at 300 kV and equipped with a Falcon-4i direct electron detector and Selectris energy filter (Thermo Fisher Scientific, Waltham, MA, USA) using 10 eV slit. The data were acquired at 165 000 x nominal magnification with a calibrated pixel size of 0.728 Å. A total of 10 065 EER formatted exposures were collected in 657 raw frames with a total dose of 28.14 e^-^/Å2 (14.06 e^-^/pixel/s) over 2.14 s with a set defocus between -0.7 to -1.3 µm.

#### Preliminary dataset1

Collected on a Glacios TEM operated at 200 kV equipped with a Falcon-III direct electron detector. The data were acquired at 150 000 x nominal magnification with a pixel size of 0.952 Å. A total of 3 490 movies were collected in 30 frames with a total dose of 30.97 e^-^/Å2 (1.01 e^-^/pixel/s) over 27.79 s with a set defocus between -0.6 to -1.4 µm.

#### Preliminary dataset2

Collected on a Titan Krios G2 operated at 300 kV and equipped with a K3 BioQuantum direct electron detector and energy filter using 20 eV slit. The data were acquired at 105 000 x nominal magnification with a calibrated pixel size of 0.8240 Å. A total of 8 533 movies were collected in 30 frames with a total dose of 28.52 e^-^/Å2 (16.1 e^-^/pixel/s) over 1 s with a set defocus between -0.8 to -1.4 µm.

### Cryo-EM data processing

All processing was done on cryoSPARC v4.4.1^31^ using a similar workflow. All the final maps were post-processed by global B-factor sharpening based on linear fit to the Guinier plot and locally low-pass filtered based on local resolution estimation (cryoSPARC local filtering). The global resolution of maps was estimated within an auto-tightened mask created by cryoSPARC. For all maps used for model refinement, the resolution within a mask generated from the model was also calculated and used as the reported resolution. These masks were created by calculating a synthetic map from the model using the molmap command in ChimeraX at 20 Å, thresholded at a level where the map covers the whole model, and then adding a 16-pixel soft edge.

#### Early dataset

The movies were motion-corrected using Patch Motion Correction, the CTFs were estimated using Patch CTF Estimation. Blob picker was used to produce an initial 3D reconstruction and produce templates for template picking. After template picking a particle stack was created by extracting a 600-pixel box around each particle. 2D classification was used to remove obvious picking artifacts (*e.g.* carbon edges), followed by a homogeneous refinement performed simultaneously with higher-order CTF estimation and per-particle defocus refinement. The resulting reconstruction was used for referenced-based motion correction and a new homogeneous refinement was performed followed by higher-order CTF estimation and per-particle defocus refinement, resulting in a final consensus reconstruction. The 70S ribosome was subtracted using a mask that covered the ribosome with an 18-pixel soft-edge. The resulting subtracted particles were then down-sampled to 128 pixels and a focused 3D classification was performed using a mask around the ribosomal A-site that covers EF-G and the expected FusB region with an 18-pixel soft-edge. The 3D classification was performed with 120 classes, while keeping the input per-particle scales. The different resulting classes were homogeneously refined using the original full-sized non-subtracted particles. The particles in the classes containing the EF-G•70S complex were down-sampled to 128 pixels and a focused classification inside a SSU-mask was performed with 20 classes, while keeping the input per-particle scales. Homogeneous refinement using the original particles resulted in the final CHI and POST structures. A schematic of the processing workflow is shown in Supplementary Figure 8.

#### 25 s dataset

Processing was done in a similar way as for the early dataset. 2D classification was followed by ab-initio and heterogeneous refinement with 5 classes to remove additional non-ribosomal particles. Focused 3D classification with an A-site mask was performed with 100 classes and using a 1% convergence criterion. The focused classification over SSU was performed with 10 classes. A schematic of the processing workflow is shown in Supplementary Figure 9.

#### FusB•Sa70S dataset

The EER movies were motion-corrected in 41 frames using Patch Motion Correction and the CTF parameters were estimated using CTFFIND4. After that, processing mostly followed the 25 s dataset. Ab-initio and heterogeneous refinement to remove non-ribosomal particles were done with 4 classes. Non-uniform refinement with higher order CTF estimation and per-particle defocus refinement was used to obtain the reference for referenced-based motion correction and then again to calculate the final consensus reconstruction. For subtracting the 70S ribosome, a mask with a 12 pixel soft-edge was used. The focused 3D classification at the A-site was done using a spherical 50 Å mask around the ribosomal A-site with a 12 pixel soft-edge and 40 classes. A schematic of the processing workflow is shown in Supplementary Figure 10.

#### Preliminary dataset1

Processing was done in a similar way as the early dataset. After movie preprocessing and blob picking, particles were extracted with 416-pixel box size binned to 128 pixels. Ab-initio reconstruction followed by heterogeneous refinement was performed with 10 classes. The ribosomal particles were re-extracted at full resolution. The consensus reconstruction was used for signal subtraction using a mask that covered the 70S ribosome with a 12-pixel soft-edge. Focused 3D classification was performed using a mask around the ribosomal A-site that covers EF-G and the expected FusB region with a 12-pixel soft-edge with 60 classes. Focused classification over SSU was performed with 5 classes. A schematic of the processing workflow is shown in Supplementary Figure 10.

#### Preliminary dataset2

Processing was done in a similar way as the early dataset. After template picking, particles were extracted with a 512-pixel box size. For focused 3D classification around the ribosomal A-site the 70S ribosome was subtracted using a mask with a 12-pixel soft-edge and using 100 classes. Focused classification over the SSU was performed with 20 classes. A schematic of the processing workflow is shown in Supplementary Figure 11.

### Model building

All models were built by first rigid-body fitting and refining each chain in the starting model using Coot^32^ against the post-processed map followed by final refinement using Servalcat^33^. Afterwards, the structures were revised manually in areas relevant to this study and validation issues were corrected. Validation was performed using Phenix^34^ and the MolProbity web server^35^. Refinement and validation information is found in Table 1.

#### FusB•Sa70S:SSU

The starting model was the 2.4 Å *S. aureus* FA-CP-locked structure (PDBID: 8P2F)^17^. For FusB, chain A from the crystal structure of FusB (4ADN)^18^ was used, and for protein bS21 an AlphaFold2^36^ prediction using ColabFold^37^ was used. Waters were added using Coot prior to the final refinement with Servalcat. The map used was from the FusB•*Sa*70S dataset.

#### FusB•Sa70S:LSU

The starting model was the refined FusB•Sa70S:SSU model. The map used was from the FusB•*Sa*70S dataset.

#### FusB•70S:SSU

The staring model consisted of PDBIDs 8CGK, 8CGJ, and 8CF1^38^. H43-44 were added from the *E. coli* FA-locked complex (PDBID 7N2C)^39^, FusB, tRNA and mRNA from *FusB•Sa70S:SSU* and remaining missing fragments from PDBID 7K00^40^. The map used was from the 25 s dataset.

#### FusB•70S:LSU

The starting model was the refined *FusB•70S:SSU* model. The map used was from the 25 s dataset.

#### FusB•EF-G•70S and FusB•EF-G•70S*

The starting model was the refined *FusB•70S:SSU* model except for FusB, and FusB•EF-G from an AlphaFold2 prediction using ColabFold. The map used was from the early dataset for FusB•EF-G•70S, and the 25 s dataset for FusB•EF-G•70S*.

#### CHI and POST

The starting model was the refined *FusB•70S:SSU* model. FusB, EF-G and tRNA came from the 2.5 Å *S. aureus* FA-locked structure (PDBID: 8P2H)^17^.The maps were from the early dataset.

## Multiple sequence alignment

The alignment was performed using Clustal Omega^41,42^ at the EMBL-EBI web server^43^ and displayed using Jalview^44^.

## Figures

Figures were rendered using UCSF ChimeraX^45^, and assembled in Affinity Designer (Serif Europe Ltd, Nottinghamsire, United Kingdom).

## Data availability

The cryo-EM maps and models in this study have been deposited in the Electron Microscopy Data Bank under the accession codes EMDB-51350 (CHI), EMDB-51351 (POST), EMDB-51352 (FusB•EF-G•70S), EMDB-51353 (FusB•EF-G•70S*), EMDB-51354 (FusB•70S:SSU), EMDB-51355 (FusB•70S:LSU), EMDB-51356 (FusB•Sa70S:SSU), EMDB-51357 (FusB•Sa70S:LSU), and Protein Data Bank under the accession codes 9GHA (CHI), 9GHB (POST), 9GHC (FusB•EF-G•70S), 9GHD (FusB•EF-G•70S*), 9GHE (FusB•70S:SSU), 9GHF (FusB•70S:LSU), 9GHG (FusB•Sa70S:SSU), 9GHH (FusB•Sa70S:LSU). Custom Python scripts used during analysis of the structures are available on Git Hub (https://github.com/adriangl97/pdb_python_tools).

## Author contributions

A.G.L. purified FusB and EF-G, prepared the cryo-EM samples, processed the cryo-EM data and modeled the structures. X.G. and S.S. performed the di and tri-peptide assays and purified the components. D.S.D.L. and M.S. supervised the structure work.

A.G.L. and M.S. wrote the manuscript with help from all the authors. M.S. and S.S. secured funding.

## Acknowledgements

We thank Magnus Johansson for comments on the manuscript and on experimental design. We acknowledge the use of the Cryo-EM Uppsala facility for grid preparation and screening, funded by the Department of Cell and Molecular Biology, the Disciplinary Domains of Science and Technology and of Medicine and Pharmacy at Uppsala University. Cryo-EM data were collected at the Cryo-EM Swedish National Facility funded by the Knut and Alice Wallenberg Foundation, the Erling-Persson Family Foundation, and the Kempe Foundation; SciLifeLab; Stockholm University; and Umeå University. This work benefited from access to Diamond Light Source and has been supported by iNEXT-Discovery project number 26535, funded by the Horizon 2020 program of the European Commission. We acknowledge access and support of the cryo-EM facilities at the UK national electron Bio-Imaging Centre (eBIC), proposal BI35989. The data processing was enabled by the Berzelius resource provided by the Knut and Alice Wallenberg Foundation at the National Supercomputer Centre. This research was funded by grants from Uppsala Antibiotic Center to M.S., from the Swedish Research Council (2017-03827 and 2022-04511 to M.S.; 2023-05237, 2018-05946 and 2018-05498 to S.S.; 2016-06264 to M.S. and S.S.) and from the Knut and Alice Wallenberg Foundation (KAW 2017.0055) to S.S. A.G.L. has received a fellowship from the Sven and Lilly Lawski Foundation.

---

**Extended data figure 1.**
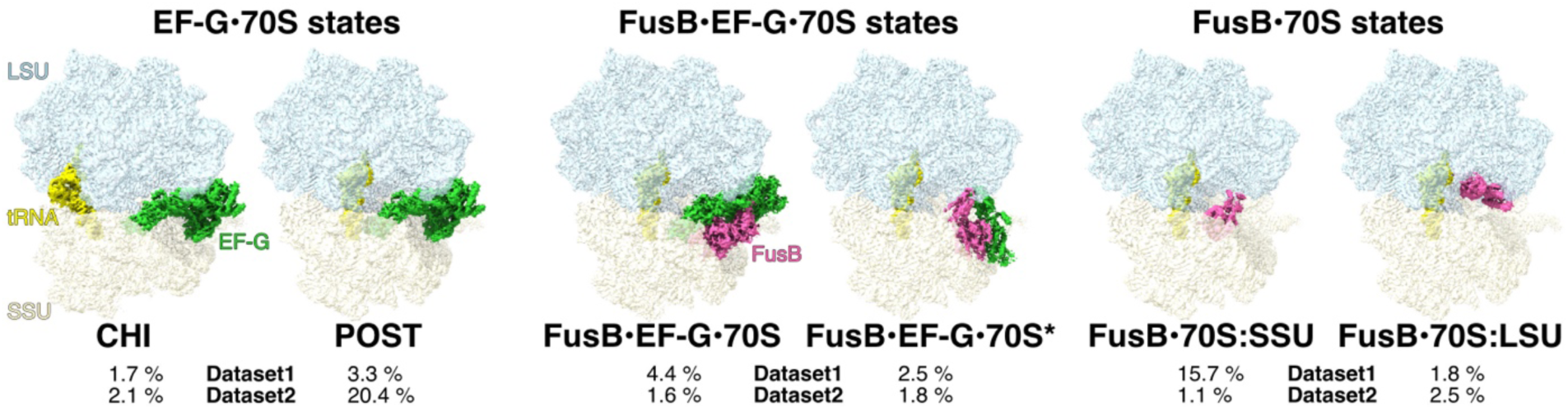
Fraction of the different states in FusB-mediated rescue in two different regions of the same grid in a preliminary experiment. This grid was prepared by mixing 1 µl of FusB into 2.5 µl of FA-complex, freezing 10 s after FusB addition. Ribosome states with empty A-site, low-occupancy EF-G or A-site tRNA correspond to the remaining particle populations (70.7 % in Dataset 1 and 73.0 % in Dataset 2).

**Extended Data Figure 2.**
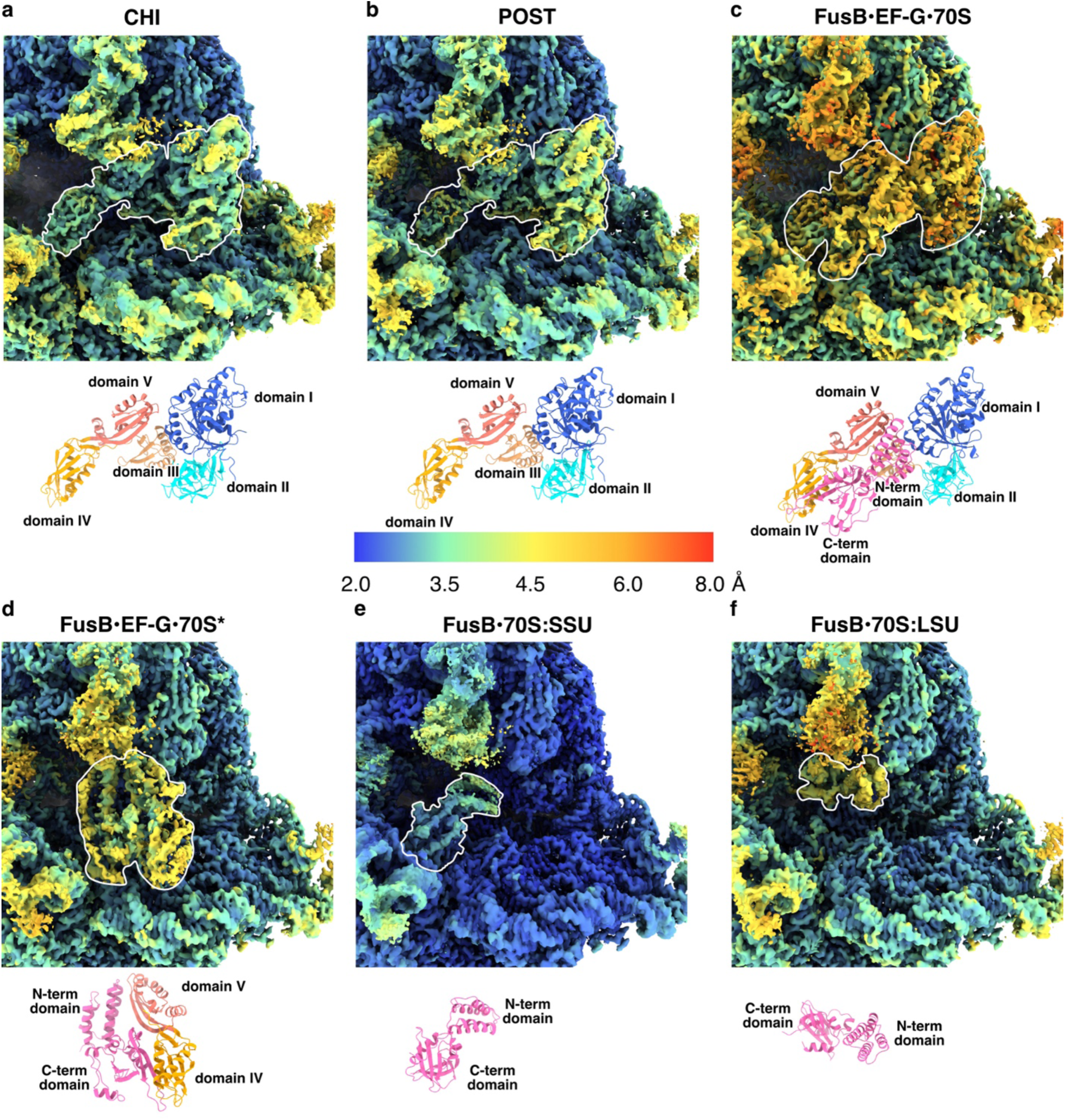
Cryo-EM maps around the ribosomal A-site from the early and 25 s datasets colored by local resolution estimation from cryoSPARC. The cartoon representation of the EF-G and FusB models in the same orientation is shown below each panel. FusB and EF-G are outlined on the maps. **(a)** CHI. **(b)** POST. **(c)** FusB•EF-G•70S. **(d)** FusB•EF-G•70S*. **(e)** FusB•70S:SSU. **(f)** FusB•70S:LSU.

