## Supplementary Figures for "Structural mechanism of FusB-mediated rescue from fusidic acid inhibition of protein synthesis"

**Content:**

**Supplementary Figures 1-12**

**Supplementary References**

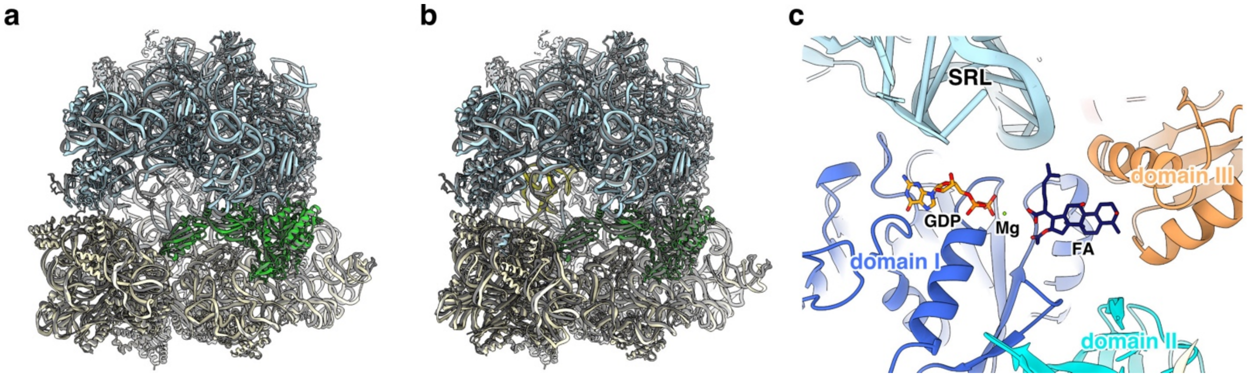


**Supplementary Figure 1. (a)** Comparison between the CHI state with *S. aureus* EF-G and *E. coli* 70S (LSU in light blue, SSU in light yellow, tRNA in yellow and EF-G in green), and *S. aureus* 70S (PDBID: 8P2H, gray)[1]. **(b)** Comparison between the POST state with *S. aureus* EF-G and *E. coli* 70S and *S. aureus* 70S (PDBID: 8P2F)[1]. Colors as in a. The structures are aligned by 23S rRNA. **(c)** FA-binding pocket in POST-state EF-G (colored by domain) with FA (dark blue), LSU (light blue), GDP (orange) and Mg (green).

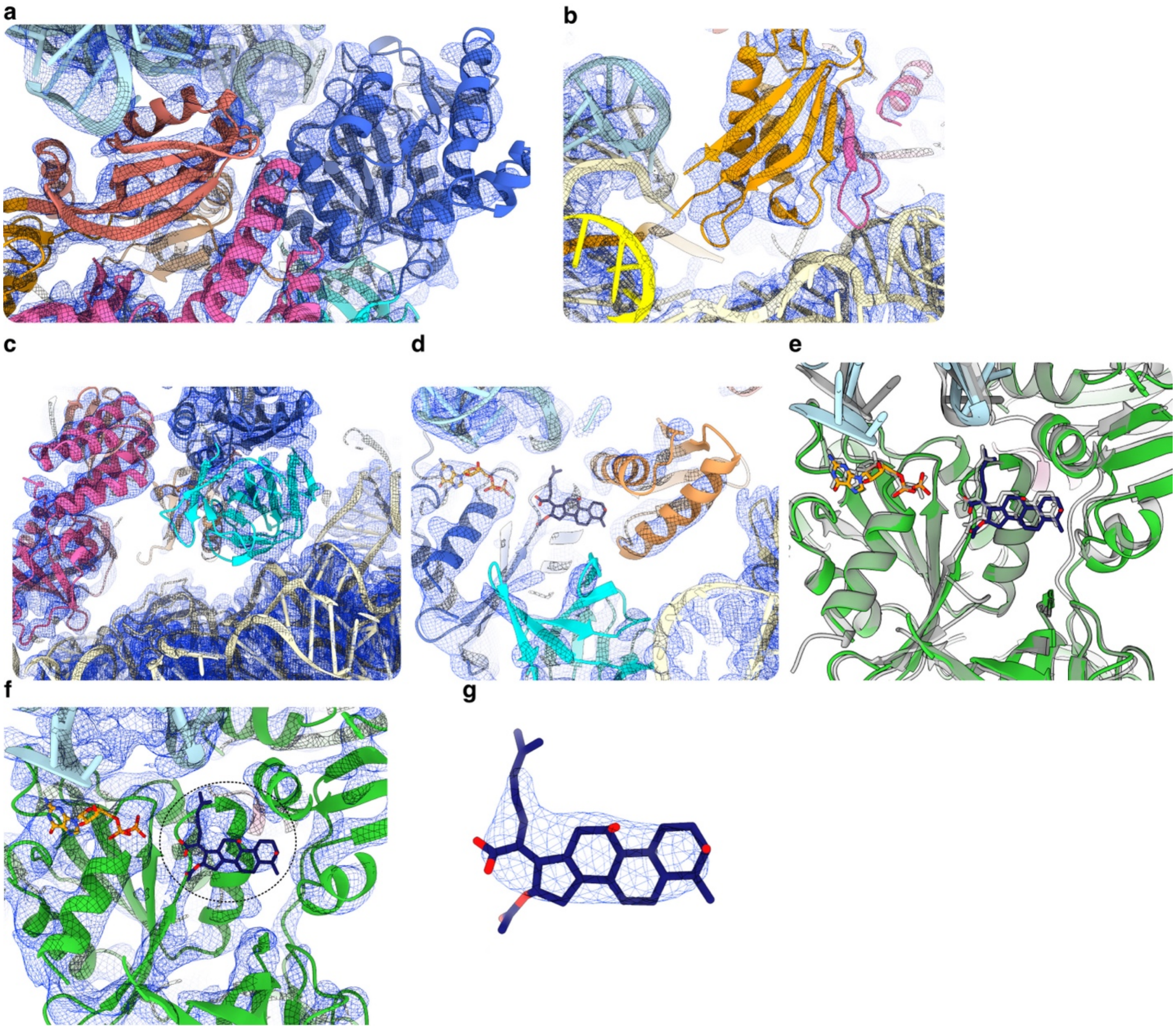


**Supplementary Figure 2.** Local filtered map of FusB•EF-G•70S around the regions shown in Figure 2d-g. **(a)** As Figure 2d. **(b)** As Figure 2e. **(c)** As Figure 2f. **(d)** As Figure 2g. **(e)** Structure alignment of domains I-II of EF-G between POST (gray) and FusB•EF-G•70S showing the FA-binding pocket, with LSU (light blue), EF-G (green), FA (dark blue), and GDP (orange). **(f)** As e, with local filtered map shown. **(g)** Segmented map of FusB•EF-G•70S  2 Å around FA.

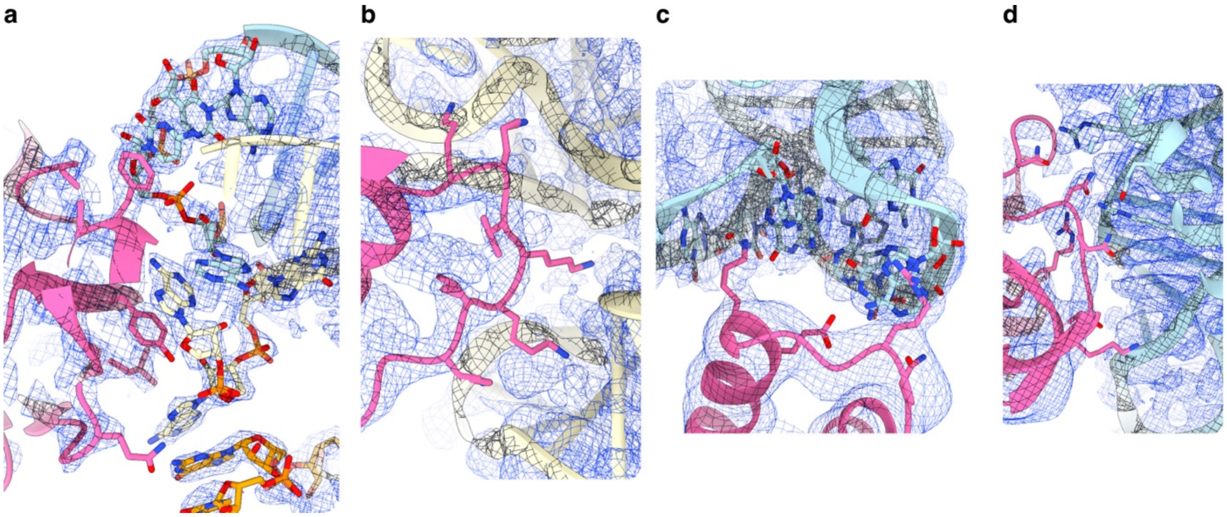


**Supplementary Figure 3.** Local filtered maps of FusB•SSU and FusB•LSU around the regions shown in Figure 3b, c, e and f. **(a)** As Figure 3b. **(b)** As Figure 3c. **(c)** As Figure 3e. **(d)** As Figure 3f.

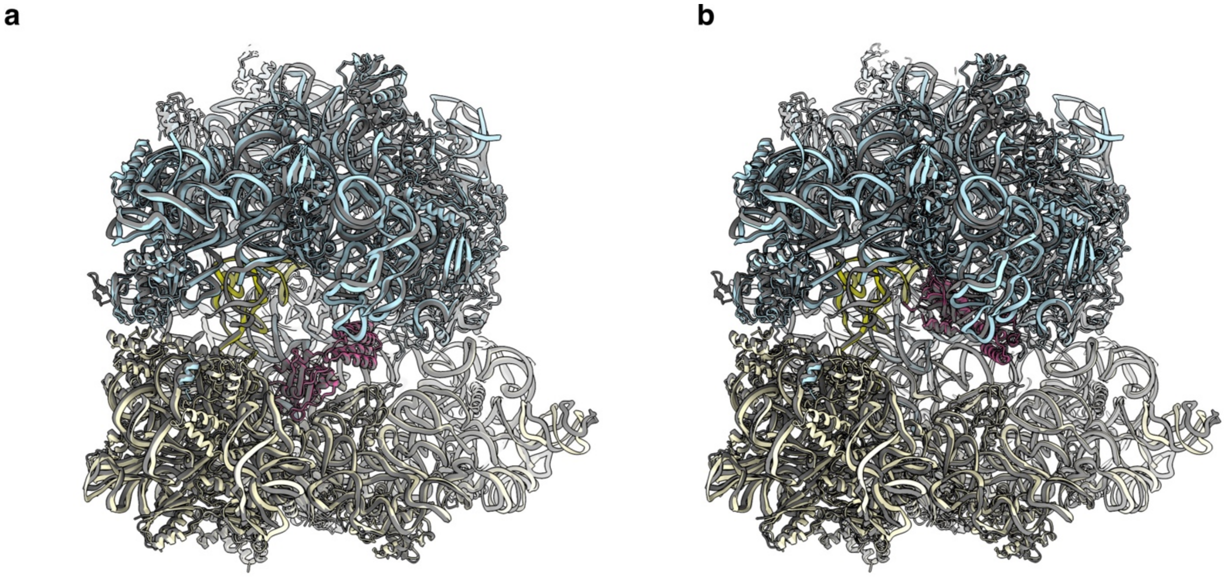


**Supplementary Figure 4.** Comparison of FusB complexes with *E. coli* and *S. aureus* 70S **(a)** Overlay of FusB•70S:SSU (LSU in light blue, SSU in light yellow, tRNA in yellow, and FusB in pink), and FusB•Sa70S:SSU (gray). **(b)** Overlay of FusB•70S:LSU (colors as in a), and FusB•Sa70S:LSU (gray).

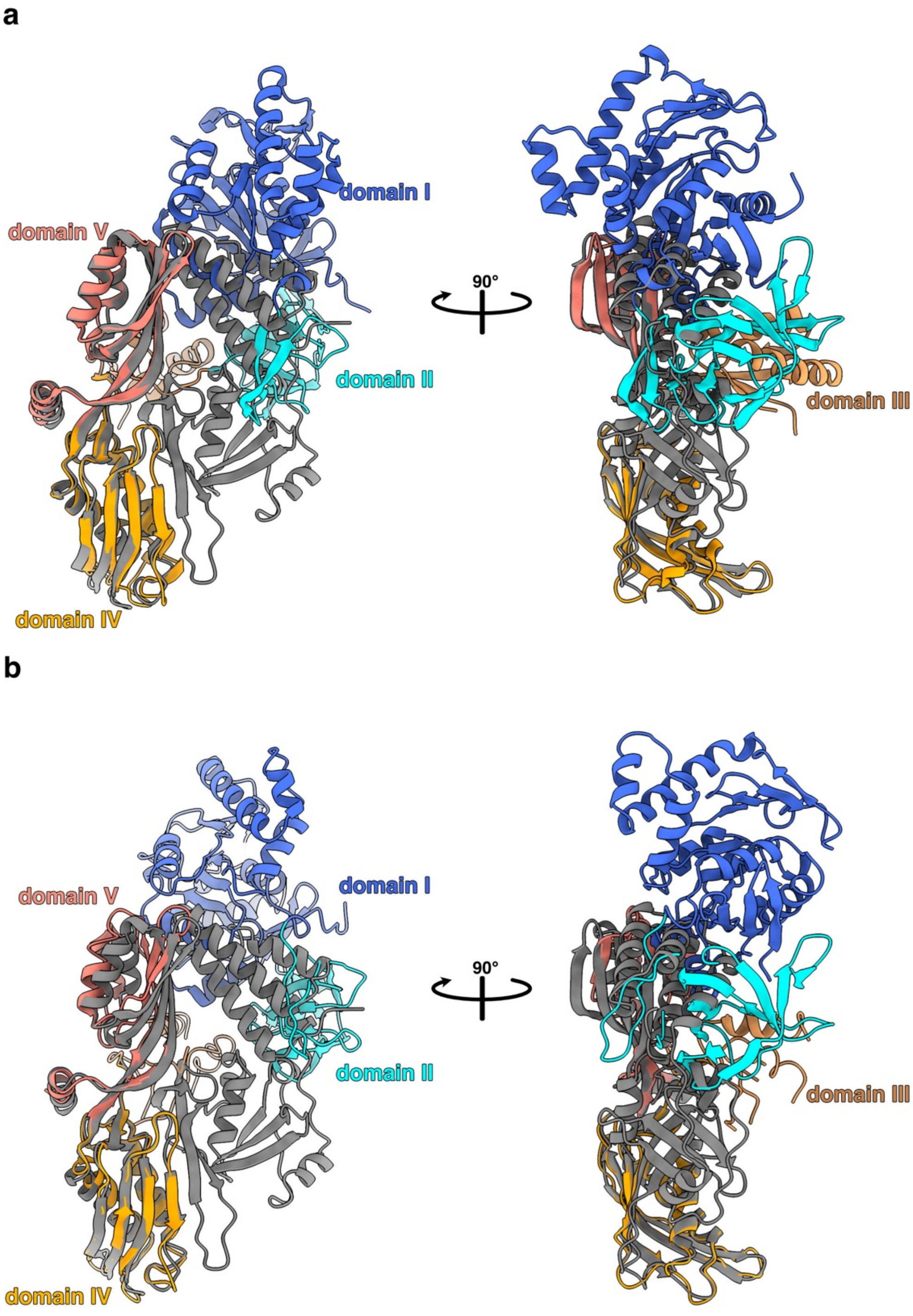


**Supplementary Figure 5.** Structure aligment based on domains IV-V of EF-G between FusB•EF-G in FusB•EF-G•70S (EF-G in green and FusB in pink) with crystal structures of EF-G shows that both domains of FusB would clash with domains I-II and the linker regions between domains II, III and IV. **(a)** Comparison between FusB•EF-G and the crystal structure of *S. aureus* EF-G (PDBID: 2XEX, colored by domain)[2]. **(b)** Comparison between FusB•EF-G and the crystal structure of *Thermus thermophilus* EF-G (PDBID: 1ELO, colored by domain)[3].

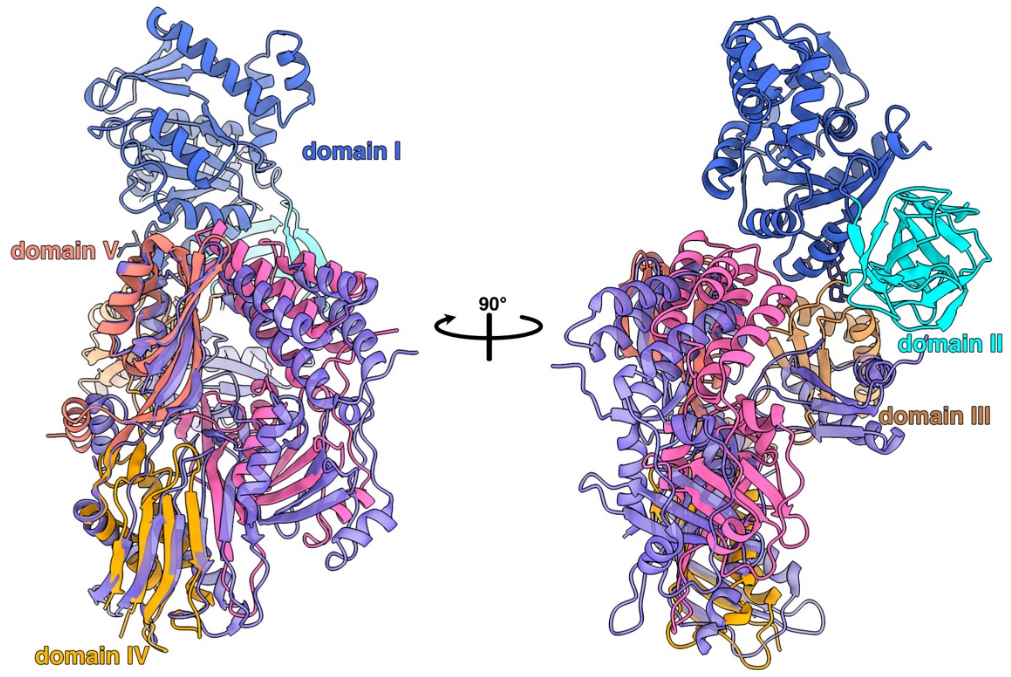


**Supplementary Figure 6.** Structure alignment based on domains IV-V of EF-G between the NMR model of FusB and EF-G domains I-III (blue, PDBID: 2MZW)[4] with FusB•EF-G in FusB•EF-G•70S (FusB in pink and EF-G colored by domain). This results in an RMSD of 14.6 Å over 213 Cα atoms for FusB.

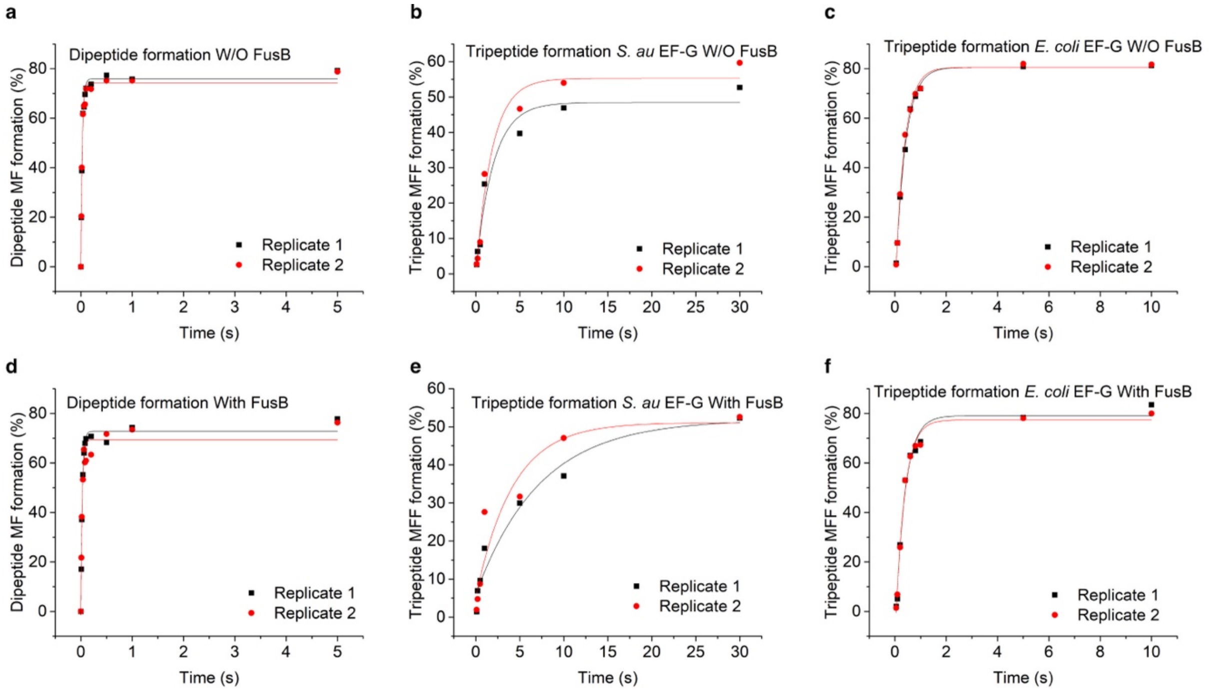


**Supplementary Figure 7.** Individual di- and tripeptide formation experiments in absence and presence of FusB (Figure 4), fitted to a single exponential curve. **(a-c)** without FusB, **(d-f)** with FusB.

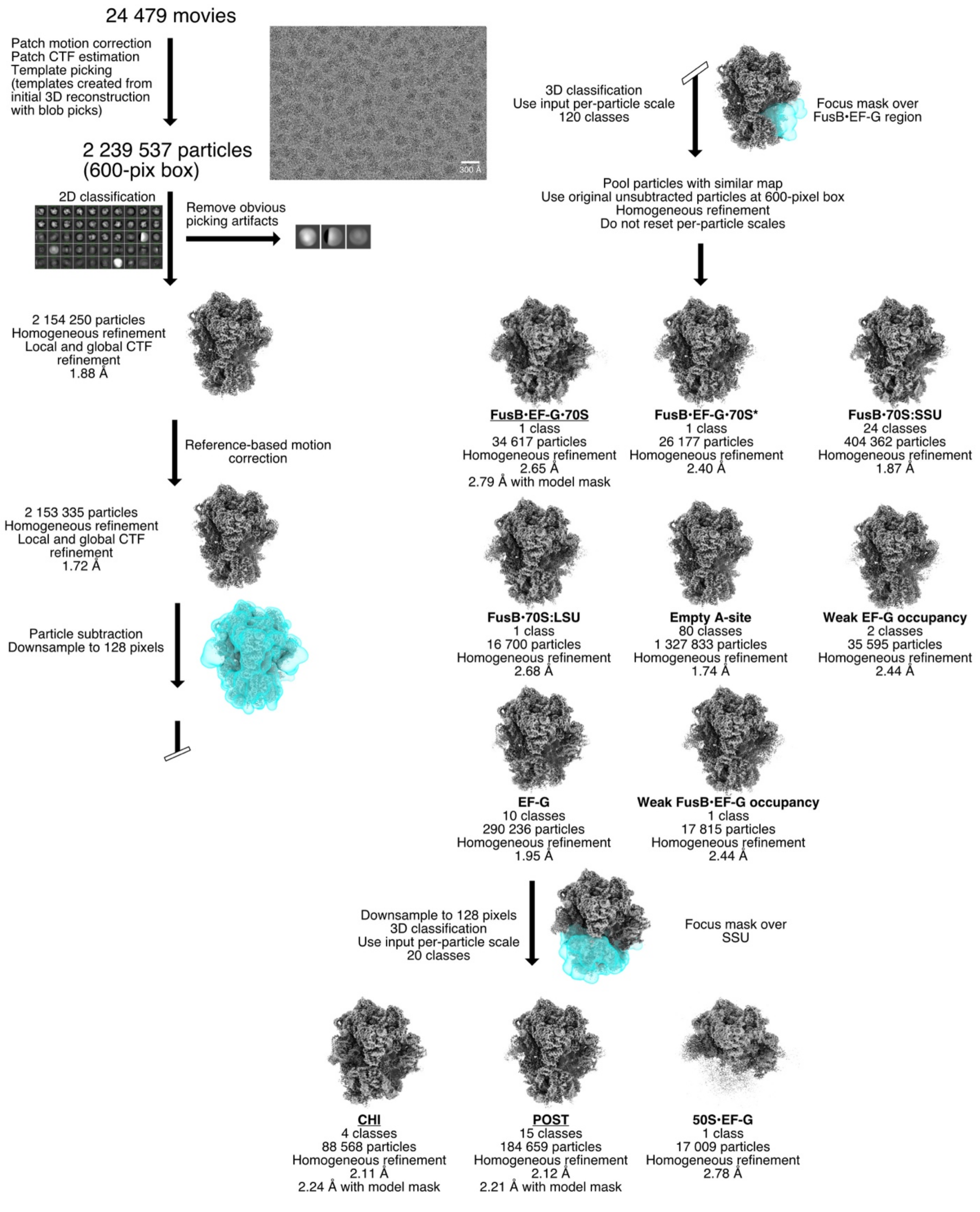


**Supplementary Figure 8.** Processing workflow for the early dataset using cryoSPARC v4.4.1. Maps used for model building and refinement are underlined.

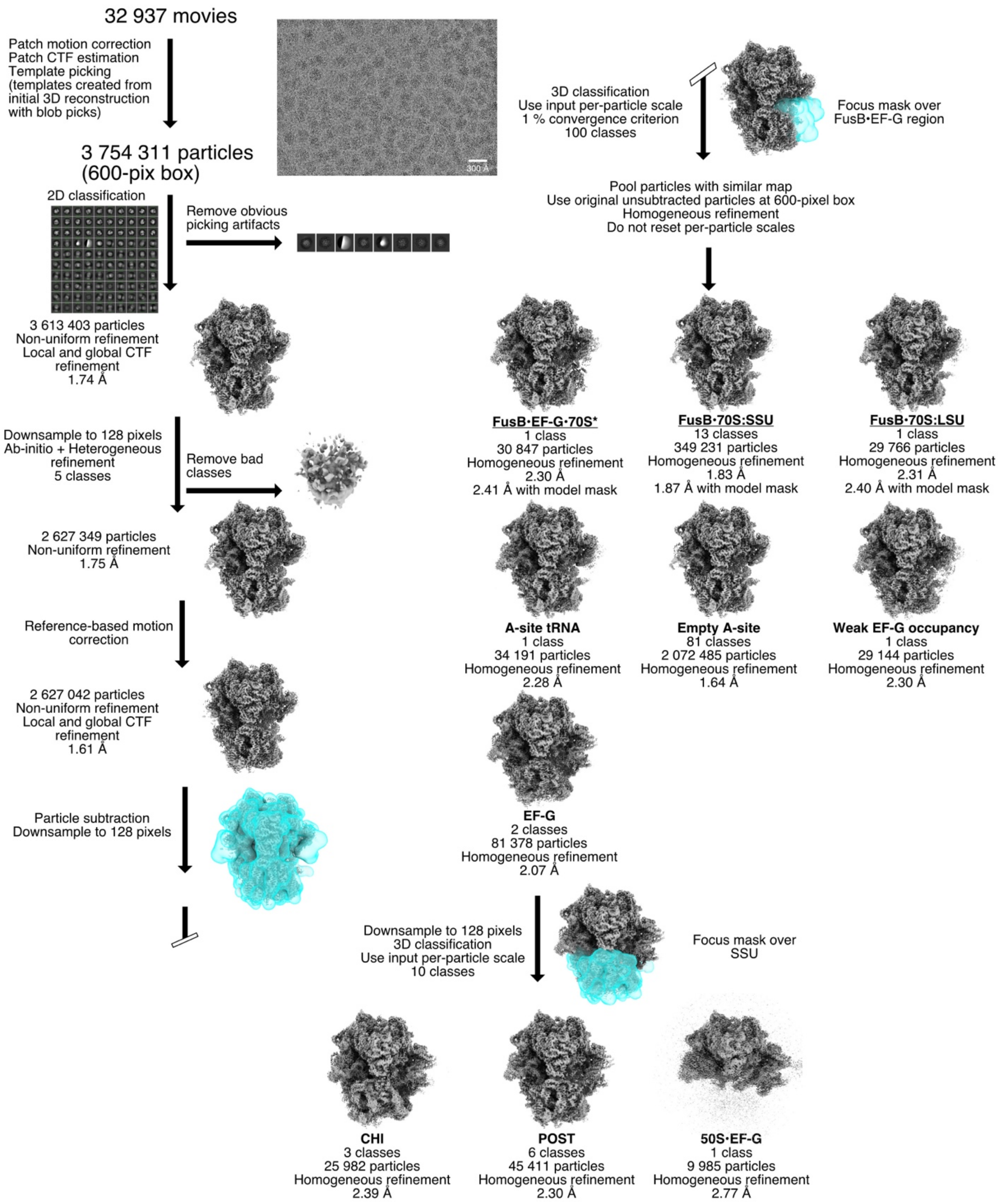


**Supplementary Figure 9.** Processing workflow for the 25 s dataset using cryoSPARC v4.4.1. Maps used for model building and refinement are underlined.

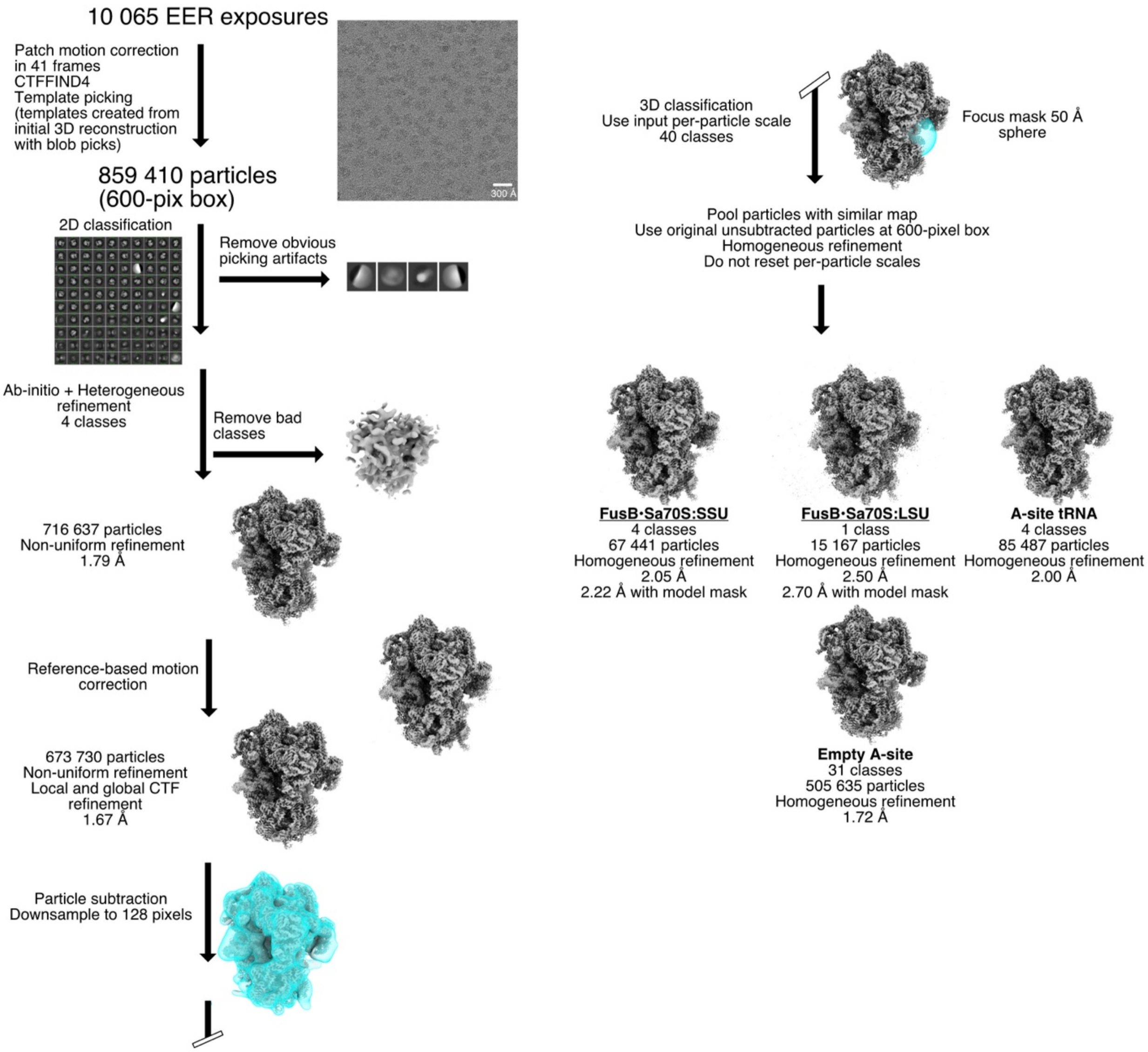


**Supplementary Figure 10.** Processing workflow for the FusB•*Sa*70S dataset using cryoSPARC v4.4.1. Maps used for model building and refinement are underlined.

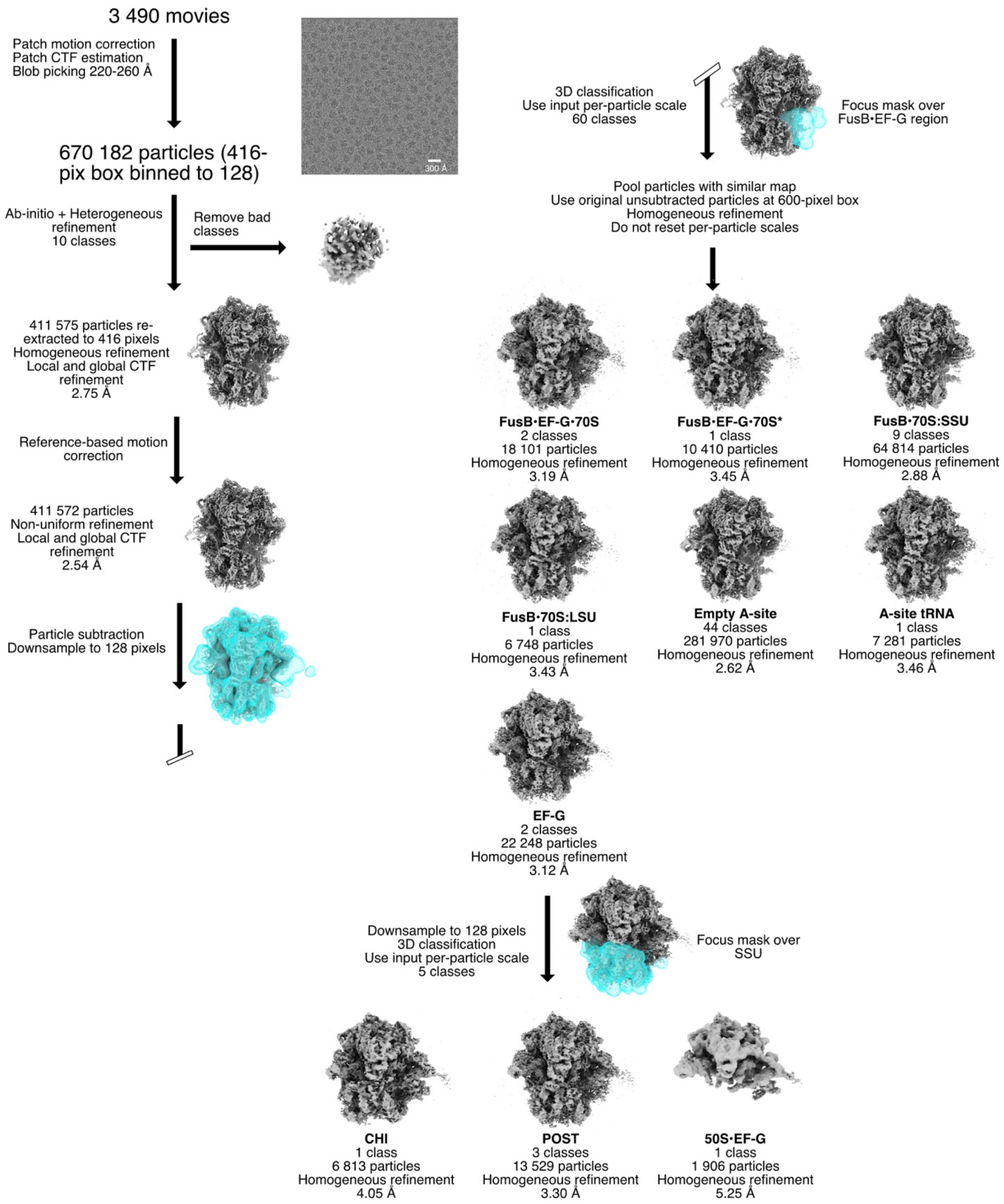


**Supplementary Figure 11.** Processing workflow for preliminary dataset1 using cryoSPARC v4.4.1.

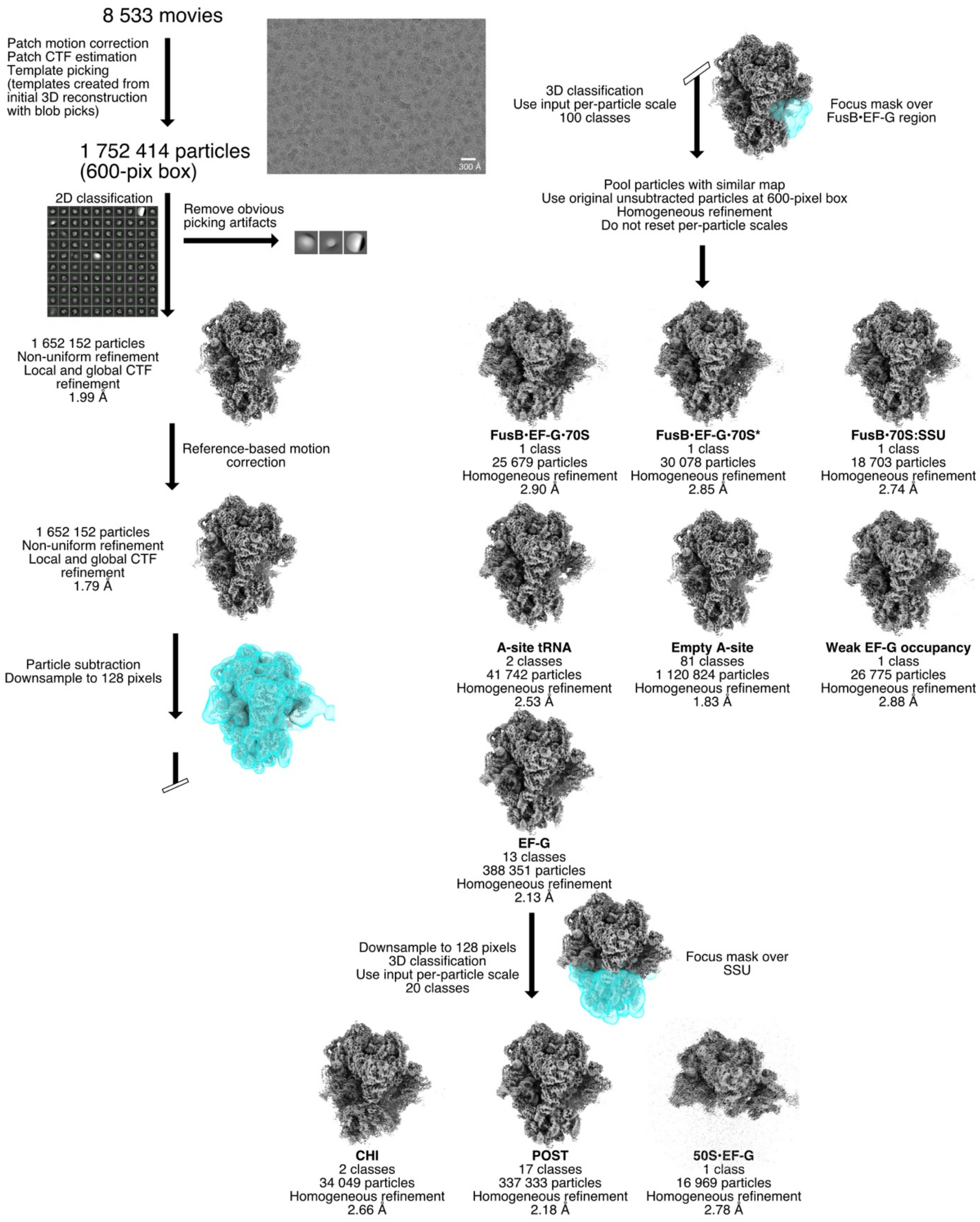


**Supplementary Figure 12.** Processing workflow for preliminary dataset2 using cryoSPARC v4.4.1.
